# KineticGP: a computational framework for genomic prediction of leaf photosynthesis traits

**DOI:** 10.1101/2025.07.16.665083

**Authors:** Rudan Xu, John Ferguson, David Hobby, Milad Rahimi-Majd, Philipp Wendering, Johannes Kromdijk, Zoran Nikoloski

## Abstract

Crop traits are the integrated outcome of genetic factors, environment effects, and their complex interactions, rendering accurate prediction from genetic markers alone a challenging problem. Here we present KineticGP, a computational framework that combines genomic prediction with genotype-specific kinetic models of C_4_ photosynthesis to make predictions of leaf photosynthesis traits across genotypes from a multiple parent advanced generation intercross maize population. Using genetic markers and gas exchange measurements from three field seasons, we show that KineticGP outperforms a baseline genomic prediction model for photosynthesis rate at saturating light by 86% for unseen genotypes across two seen seasons. In addition, KineticGP allowed surveying the genetic variability in enzyme kinetic parameters that can be used to raise targets for improvement of photosynthesis. The approach paves the way for interrogating and integrating the dynamic interactions between genotype and environment to improve the prediction accuracy of photosynthetic traits.

## Introduction

Genomic prediction (GP) has revolutionized plant breeding by significantly reducing the time for the development of genotypes with desired traits (Massman et al., 2013; Vivek et al., 2017). GP relies on building and utilizing machine learning models for traits of interest using genetic markers, from a population of genotypes, as predictors. GP has been shown to successfully predict yield-related traits for major crops, including maize (Zhao et al., 2012), barley (Lorenz et al., 2012), and wheat (Rutkoski et al., 2012).

The key challenge of GP is the generalizability of the machine learning models to genotypes and environments not seen in the process of model training (Desta & Ortiz, 2014; Heslot et al., 2015; Voss-Fels et al., 2019). This challenge is largely due to the presence of genotype-by-environment (G×E) interactions, whereby genotypes may respond differently to environmental changes, prominent for the majority of yield-related traits (de Leon et al., 2016; Kang, 2004). While advances in quantitative genetics have resulted in GP models that consider G×E interactions (Des Marais et al., 2013; Malosetti et al., 2013), the generalizability of these models to fully unseen conditions, typical for future climate scenarios, remains poorly explored. The reason for lack of generalizability is due to the purely statistical nature of the underlying models, that fail to capture the intricate way in which molecular processes respond to environmental changes.

To better incorporate effects of different environments, hybrid models represent an alternative to the purely machine learning models in GP. This strategy combines machine learning approaches with mechanistic models of the underlying biological processes that determine the traits of interest. For instance, metabolic models (Tong et al., 2020) or crop growth models (Messina et al., 2018) have been combined with GP to predict rosette growth in *Arabidopsis thaliana*and crop yield in *Zea mays*, respectively. In the example of *A. thaliana*, genotype-specific metabolic models were used to predict steady-state fluxes for which GP models were trained; the statistical models were in turn used together with the metabolic models to predict growth for environments that were not encountered in model training, but could be simulated by the mechanistic model. Similarly, crop growth models for maize rely on large multi-environment data sets to estimate model parameters that are in turn predicted by genetic markers and used in simulations of unseen environments (Messina et al., 2018). Despite these advances, the use of GP with mechanistic models is still regarded as a conceptual advance whose potential for practical applications necessitates further rigorous testing (Onogi, 2022).

Here, we focus on photosynthesis because its efficiency in field experiments is considerably below the theoretical upper limit (Zhu et al., 2010); moreover, many studies have shown that bioengineering of the underlying biological processes, such as: light induction of the Calvin cycle, photoprotection, and photorespiration, could improve photosynthetic efficiency and crop yield (Burgess et al., 2023; Kromdijk et al., 2016; Sales et al., 2021). Application of the hybrid modeling frameworks to improving photosynthetic efficiency has not yet been attempted; this hybrid framework can, in principle, rely on simplified steady-state models (Farquhar et al., 1980; Von Caemmerer, 2000), that appear in crop growth models (He et al., 2024), as well as more elaborate kinetic models of photosynthesis (Arnold & Nikoloski, 2011; Wang et al., 2021; Zhu et al., 2013).

Steady-state models of photosynthesis have been applied to estimate photosynthesis-related traits, but are limited to surveying the genetic variability in the efficiency and maximal velocity of enzymes beyond RuBisCO (Farquhar et al., 1980; Von Caemmerer, 2000). Application of more elaborate kinetic models necessitates extensive estimation of model parameters using genotype-specific data of photosynthesis-related traits. However, once calibrated for a population of genotypes, kinetic models can be readily applied to simulate plant responses to unmeasured environmental fluctuations, such as sudden changes in light intensity and temperature, thereby capturing G×E interactions.

Here, to enable the prediction of leaf photosynthesis in future climate scenarios, we first estimate genotype-specific kinetic parameters using an existing kinetic model of C_4_ photosynthesis (Wang et al., 2021) and gas exchange measurements from a multiple parent advanced generation intercross (MAGIC) maize population. The resulting approach, called KineticGP, integrates genotype-specific kinetic model parameterization with genomic prediction of kinetic parameters and simulation of leaf photosynthesis. Using KineticGP, we showed that photosynthesis-related traits can be predicted for unseen genotypes, with prediction performance improving from 0.14 to 0.26 when using best linear unbiased predictors (BLUPs) across two seasons. KineticGP does not only identify genotypes with improved photosynthetic efficiency in unseen environments, but also provides mechanistic insight in enzyme parameters that affect photosynthesis.

## Results and Discussion

### Default kinetic model fails to reproduce measured net photosynthesis

To demonstrate the KineticGP proof-of-concept, we gathered light and CO_2_ response gas exchange measurements for 68 genotypes of a maize MAGIC population, grown in two field seasons in 2022 and 2023. These data included photosynthesis rate (*A*) and stomatal conductance (*g_s_*) under changing environmental net CO_2_ concentration (C*_a_*), denoted by *A*-*C_a_* and *g_s_*-*C_a_* curves, respectively, as well as *A* under changing photosynthetically active radiation (*PAR*), denoted by *A*-*PAR* curve (Figure 1a, Methods).

**Figure 1:**
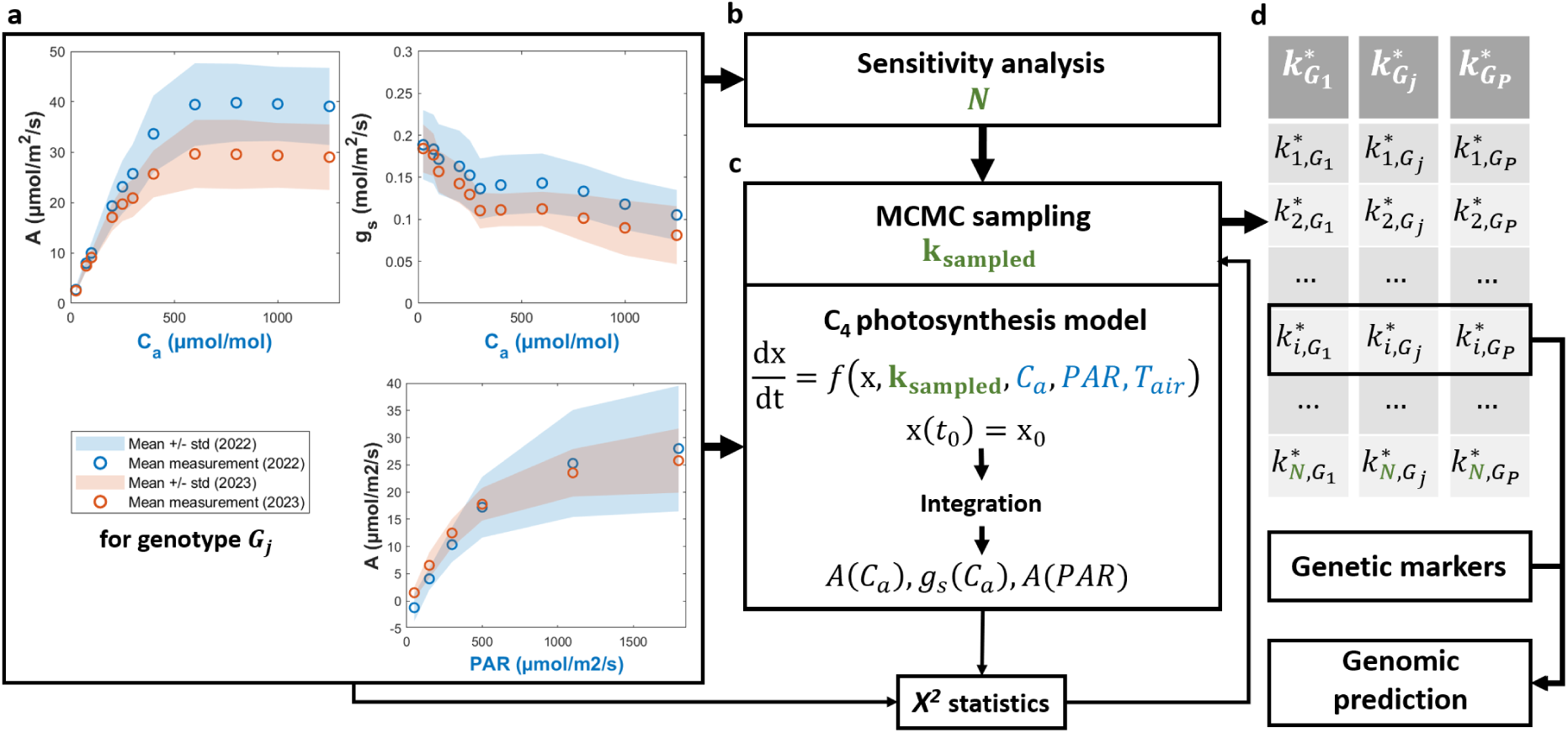
Illustration of the KineticGP framework for prediction of photosynthesis-related traits. (a) Gas exchange measurements of photosynthetic rate (*A*) and stomatal conductance (*g_s_*) across varying ambient CO_2_ concentrations (*C_a_*), as well as photosynthetic rate (*A*) under different levels of photosynthetically active radiation (*PAR*) for a given genotype (*G_j_*), were used as input data. Circles indicate the mean values across samples, and the shaded areas represent the standard deviation around the mean. The measured *C_a_* and *PAR* levels were then used to simulate the photosynthetic rate (*A*). (b) Sensitivity analysis was conducted for each kinetic parameter individually across all genotypes, and the resulting control coefficients were then used to identify which parameters should be jointly estimated (*N* parameters). (c) Markov chain Monte Carlo (MCMC) was used to estimate the kinetic parameters. The process began with the default parameter values and iteratively sampled new parameter sets (*k_sampled_*). After each sampling step, the C_4_ photosynthesis model (Wang et al., 2021) was simulated at the measured *C_a_*and *PAR* levels from (a) to generate *A*(*C_a_*), *g_s_*(*C_a_*), and *A*(*PAR*) for both seasons. The goodness-of-fit, quantified by the *χ*^2^ statistic comparing measured and simulated *A* and *g_s_*, guided the parameter estimation. (d) This procedure yielded the optimal parameter values for each genotype *G_j_* (*k_G_j__^*^*). The estimation was repeated for all genotypes (*G_p_* denotes the total number of genotypes), and the resulting estimated parameters were then used as training responses for genomic prediction (GP) based on genetic markers.

The experimental data revealed significant variation in net photosynthesis across genotypes, with the lowest *A*-*C_a_* plateau measured at 20.43 and 16.29 µmol m^-2^ s^-1^ in the 2022 and 2023 seasons, respectively, while the highest plateaus were at 52.28 and 50.05 µmol m^-2^ s^-1^ (Figure S1). In 2022, measured *A* at saturating light ranged from 9.57 to 41.45 µmol m^-2^ s^-1^, whilst in 2023, it ranged from 8.58 to 37.20 µmol m^-2^ s^-1^.

The kinetic model of *C*_4_ photosynthesis (NADP-ME-type) was used to simulate leaf photosynthesis in maize (Wang et al., 2021). The kinetic parameters consider key properties of enzymes, *e.g.,* maximal velocities (V_max_), and of enzyme-metabolite pairs, *e.g.*, Michaelis-Menten constants (K_m_) and inhibitor constants (K_i_). Furthermore, the model also includes activation rate constants for light-regulated enzymes and membrane permeability coefficient of metabolites. The 236 kinetic parameters can be partitioned into 164 Michaelis-Menten constants, 53 maximal velocities, and 19 additional parameters. Among the provided default parameters, Wang et al. (2021) adjusted 11 species-specific parameter values for maize, sorghum, and sugarcane.

First, we assessed the simulated photosynthetic rate and stomatal conductance under varying *C_a_* and *PAR* using the default maize-specific kinetic parameters. The resulting simulation of CO_2_ response showed a rapid increase in photosynthetic rate at low *C_a_* levels, with the plateau occurring around 400 µmol CO_2_ mol^-1^ and maximum rate around 45 µmol m^-2^ s^-1^ (Figure S2). However, experimental data indicated that the plateau typically occurred around *C_a_* of 800 µmol CO_2_ mol^-1^. This earlier plateau in *A*-*C_a_* curves may have resulted from overestimation of stomatal conductance, reducing diffusional limitation to negligible level, typical for non-stressed plants grown under control conditions.

The mismatch between simulation using default parameters and the measured profiles indicated the need to estimate genotype-specific kinetic parameters to achieve accurate model performance across the 68 genotypes. Furthermore, due to the large number of parameters, one way to address this issue is to first identify the parameters that most strongly influence the model fit to the experimental data.

### Sensitivity analysis reveals kinetic parameters with the largest effect on model fit

To evaluate the individual contribution of each kinetic parameter to the model’s ability to reproduce gas exchange data, we performed single-parameter optimization following the workflow in Figure 1c (*i.e*, fitting with *N* = 1). Given that gas exchange measurements were obtained across two growing seasons, potentially affecting the protein abundances, we allowed the *V_max_* parameters to vary by season. The sensitivity of the model fit to each parameter was quantified using a control coefficient, defined as the absolute value of the ratio between the logarithm of the optimized-to-initial goodness-of-fit and the logarithm of the optimized-to-initial parameter values (See Methods). Ranking the parameters by their control coefficient revealed those with the largest impact on model performance (Figure 1b).

To examine the relationship between effects of parameters and photosynthetic responses, the 68 genotypes were clustered into four groups using k-means clustering (k = 4) based on their *A*-*C_a_* and *A*-*PAR* response profiles across two seasons (Figure S3a). One cluster represented genotypes with consistently high performance across both seasons (green cluster), while another showed consistently low performance (pink cluster).

The remaining two clusters exhibited variable responses. Application of t-Distributed Stochastic Neighbor Embedding (t-SNE) with the full set of individually fitted parameters could not separate the clusters obtained based on the photosynthetic responses (Figure S3b). However, applying t-SNE to the top 40 parameters with largest control coefficients revealed a clear grouping of the high-performing genotypes (cluster of green points, Figure S3c), suggesting that a subset of kinetic parameters with the highest control coefficient is strongly associated with photosynthetic performance.

Among the top-ranked parameters, several parameters were previously estimated for maize by Wang et al. (2021): the slope of Ball–Berry model for determining the steady state stomatal conductance (*BBslope*), mitochondrial respiration (*MRd*), maximum phosphoenolpyruvate (PEP) carboxylation activity (*V_m_*-PEPC), maximum RuBisCO activity (*V_m_*-RuBisCO), and the time constant of RuBisCO activation (*τ_ActRubisco_*).

We simulated *A*-*C_a_* and *A*-*PAR* curves using individually estimated kinetic parameter and categorized the parameters based on their influence on the resulting curve shapes (Figure S4). Given that the standard deviation of measured *A* values was smaller at lower *C_a_* levels compared to higher *C_a_*, the algorithm prioritized fitting the initial slope before adjusting to the plateau. Some parameters, including *BBslope*, K_m_ of HCO_3_^-^ and of PEP for PEP carboxylase, mesophyll conductance (gm), K_m_ of CO_2_ for carbonic anhydrase (CA), improved the slope at low *C_a_* while preserving high plateau values. Other parameters, such as V_max_ of PEP carboxylase, ferredoxin-NADP+ reductase (FNR), NADP+ dependent malate dehydrogenase (MDH), and NADP+-dependent malic enzyme (ME), reduced the slope but also lowered the plateau. A third group, primarily composed of K_m_parameters, failed to improve either the slope or plateau.

For the *A*-*PAR* curves, which already fit relatively well with default parameters, individual parameters such as K_m_ of HCO_3_^-^ and PEP for PEP carboxylase and permeability coefficient for CO_2_ diffusion between mesophyll and bundle sheath cells (Perm-CO_2_), improved the match to median observed values across genotypes. However, these parameters also showed limited variation across genotypes, as reflected by narrow inter-quartile ranges (Figure S5). Other parameters led to large deviations from the measured profiles, likely due to prioritizing the fit to the *A*-*C_a_* curves at the expense of *A*-*PAR* accuracy.

Despite improvements over simulations using default parameters, simulations using individually optimized parameters still failed to achieve statistically significant fits. The reduced *χ*^2^ (see Methods for calculation) remained above one for all parameters (Figure S6, right panel), indicating poor fit.

This sensitivity analysis revealed that a subset of kinetic parameters, particularly those related to carboxylation, electron transport, and stomatal regulation, showed relatively higher influence on model fit and were strongly associated with genotype-level differences in photosynthetic performance. Whilst fitting individual parameters revealed valuable physiological insights, this procedure was insufficient to achieve statistically acceptable fits. These findings underscore the necessity of simultaneously estimating multiple key parameters to accurately capture genotype-specific gas exchange responses.

### Joint estimation of key kinetic parameters allows accurate simulation of photosynthetic profiles

The top 10, 20, 30, and 40 parameters with the highest median control coefficients across genotypes were selected for model parameterization (Figure 1c). When only one season of a *V_max_* parameter was among the top contributors, its corresponding value from the other season was also included to ensure consistency. The combination of seasons resulted in parameter subsets of 11, 20, 34, and 44 kinetic parameters for joint estimation. As in previous analyses, *V_max_* values were allowed to vary between seasons, while all other parameters were held constant across seasons but allowed to differ between genotypes (see Methods). Equilibrium constants (K_eq_), determined by thermodynamic principles, were kept constant across all genotypes and seasons and thus excluded from the parameterization. Note that the K_eq_ were updated from the original values by Wang et al. (2021), as described in Supplementary Information A.1.

Following the workflow illustrated in (Figure 1), model parameters were estimated using experimental data from both seasons for each genotype. The parameter sets that yielded reduced *χ*^2^ closest to one were selected as the final estimates, to prevent overfitting. When estimating the top 11 kinetic parameters across 68 genotypes, the median contributions to the reduced *χ*^2^ statistic were 0.22 for *A*-*C_a_*, 0.13 for *g_s_*-*C_a_* and 0.08 for *A*-*PAR* curves from 2022 (Figure 2b); comparable results were obtained for season 2023. The median total reduced *χ*^2^ statistic over the two seasons reached a value of 0.997, indicating statistically acceptable fits.

**Figure 2:**
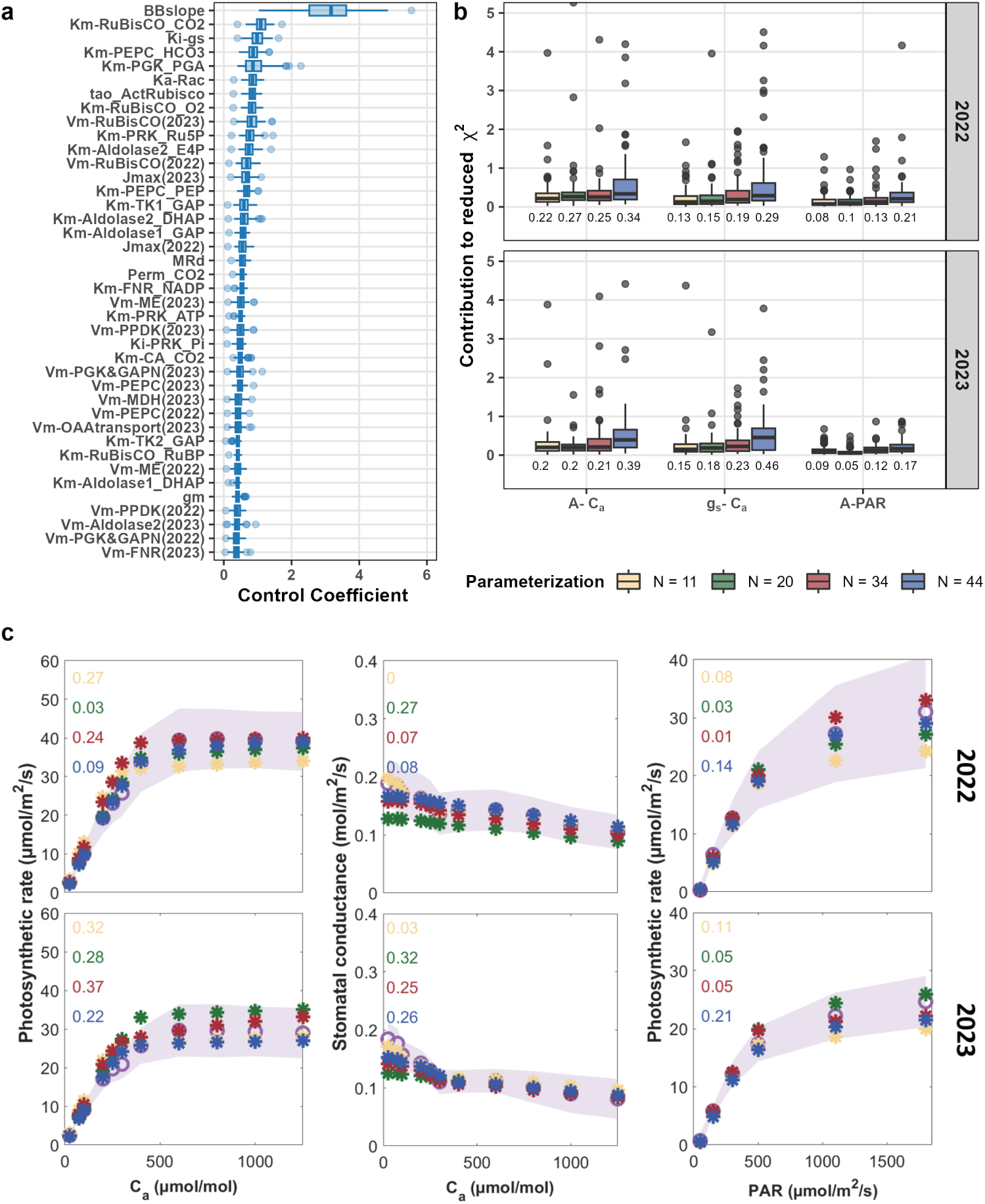
Goodness-of-fit for genotype-specific photosynthetic rate and stomatal conductance. (a) Boxplot of the top 40 kinetic parameters with respect to their control coefficients across 68 genotypes. (b) Boxplot of reduced *χ*^2^ values of the canonical curves representing photosynthetic rate and stomatal conductance over varying ambient CO_2_ levels (*A*-vs-*C_a_*and *g_s_*-vs-*C_a_* curves) and photosynthetic rate over varying photosynthetically active radiation (*A*-vs-*PAR* curve) across the 68 genotypes. The colors represent the estimation of different number of kinetic parameters-top 11 parameters (yellow), 20 parameters (green), 34 parameters (red) and 44 parameters (blue) with highest control coefficients. The reduced *χ*^2^ was calculated by dividing the *χ*^2^ values with the degrees of freedom given by the difference between number of data points and the number of estimated parameters: 45 (11 parameters), 36 (20 parameters), 22 (34 parameters) and 12 (44 parameters). (c) Examples of fits for photosynthetic response curves for a given genotype, SSA00067, in two seasons; the value of the reduced *χ*^2^ statistic of each fit is shown in each subplot. Purple circles represent the mean measured values across samples while the shaded area represent the standard deviation. Simulated values are marked with stars, colored according to the number of jointly estimated parameters as in (b).

As the number of parameters increased, the absolute model fit, as expected, improved, at the cost of stricter statistical thresholds due to the fewer degrees of freedom (Figure S7 and 2c, illustrated using genotype SSA00067 as example). As results, the median total reduced *χ*^2^ values were 0.996, 1.264, and 2.263 for the top 20, 34, and 44 parameter subsets, respectively. Across all four estimations with the different numbers of parameters, SSA00003, SSA00179, SSA00024, SSA00096, and SSA00299 showed the best overall *χ*^2^ statistic across all three canonical curves and both seasons included. In contrast, those with the highest *χ*^2^ values, indicating worse fits, included: SSA00033, SSA00075, SSA00008, SSA00367, and SSA00229 (see Figure S8-S13). Larger *χ*^2^ values were often linked to low standard deviations within genotypes, particularly for the *A*–*C_a_* and *A*–*PAR* curves.

The optimized parameter sets successfully addressed limitations of using default kinetic parameters in simulating the dynamic range of the measured photosynthetic rates. These parameters enabled accurate simulation of both the highest plateaus in *A*-*C_a_* curves (*e.g.*, SSA00239 and SSA00004) and the lowest plateaus (*e.g.*, SSA00006 and SSA00138) across two seasons. These results underscore the flexibility of the kinetic model in capturing diverse gas exchange responses across genotypes.

### Estimated kinetic parameters exhibit genetic variability and are partly heritable

The estimated values for the kinetic parameters can be used in downstream GP if they: (i) are precise (*i.e.*, show small variability) in single genotype, (ii) show variability across genotypes, and (iii) are heritable. Given only the top 11 and 20 kinetic parameters with the highest control coefficient led to significant fits, we focus on these two cases for the downstream analysis.

The precision of the estimated value for each parameter in a genotype was assessed by conducting a robustness analysis. To this end, we sampled parameter values around the optimal fit using Markov Chain Monte Carlo and constructed confidence intervals (CIs) for each parameter in the investigated genotype. The ratio between the maximum and minimum values within the 80% CI was calculated as an indicator of parameter identifiability. A parameter was considered well-determined by the photosynthetic response curves if this ratio falls between 1 and 2. When estimating 11 kinetic parameters, five of the parameters were well-determined in at least 80% of genotypes. This number increased to 18 of parameters when estimating 20 parameters (Figure S14).

To determine genetic variability of the estimated values for a single parameter, we calculated the coefficient of variation over the genotypes. Among the top 11 kinetic parameters with the highest control coefficient, bicarbonate affinity for PEP carboxylase and *BBslope* showed the smallest coefficient of variations (CVs), suggesting high conservation across genotypes. In contrast, CO_2_ affinity for RuBisCO and the light activation constant for RuBisCO activase (*K_a_R_ac_*) exhibited the highest variability (Figure 3a). For the top 20 parameters, *MRd* showed relatively small overall CVs, indicating that this parameter varies little across genotypes; by contrast, the affinities of dihydroxyacetone phosphate (DHAP) and D-erythrose-4-phosphate (E4P) to aldolase were among the most variable. The average CVs across all parameters was 75% and 81% for estimation of 11- and 20-parameters sets, respectively.

**Figure 3:**
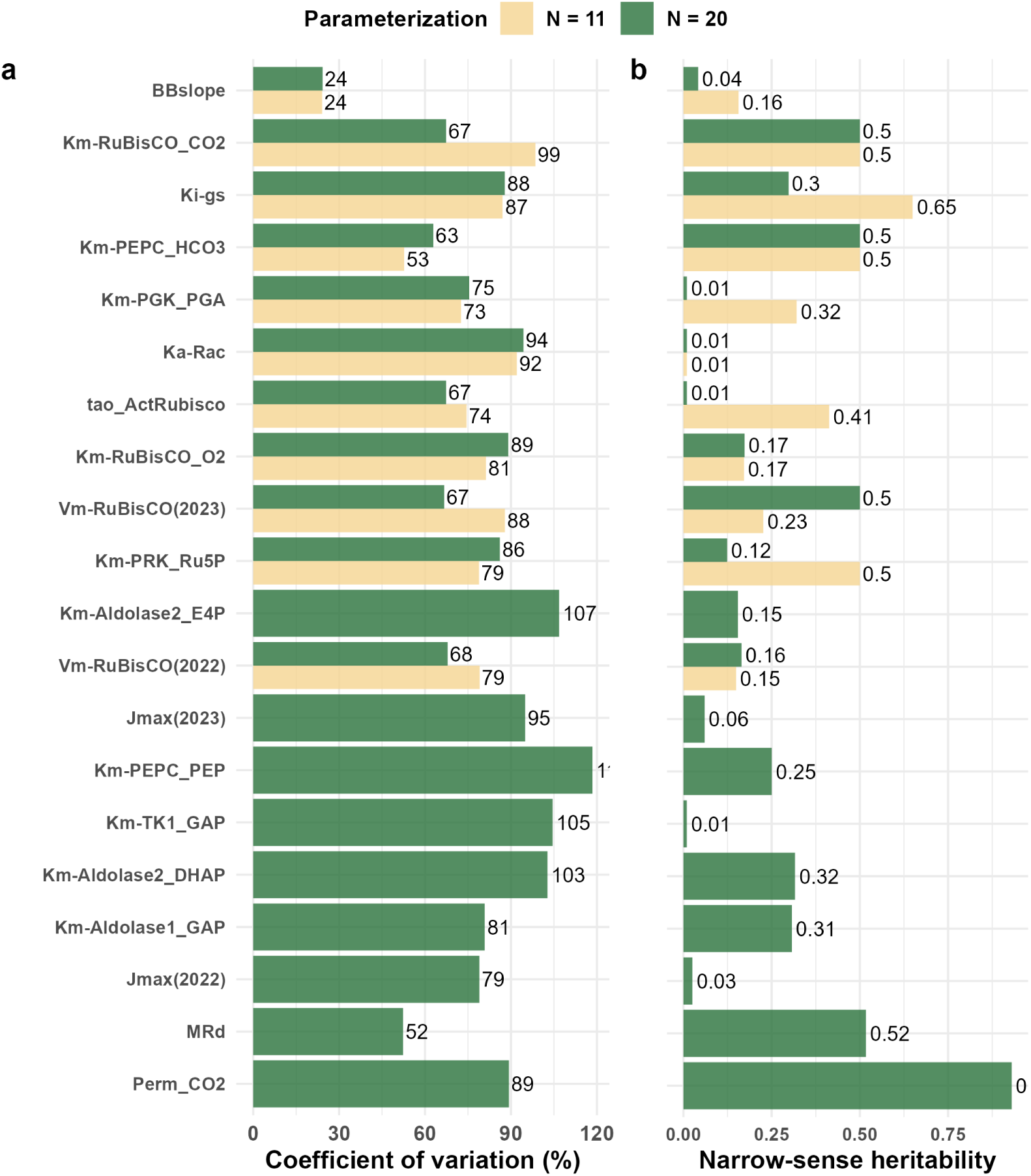
Between-genotype variation and heritability of estimated kinetic parameters. (a) Coefficient of variation (in percent) of the estimated kinetic parameters across 68 genotypes. The colors represent the estimations based on different number of kinetic parameters-top 11 parameters (yellow) and 20 parameters (green) with the highest control coefficient. (b) Narrow-sense (SNPs-based) heritability of estimated kinetic parameters.

We further calculated the marker-based (narrow-sense) heritability for each model parameter to assess their suitability for GP. Among the estimated top 11 parameters, six showed heritability values greater than 0.3, whilst the estimation of 20 parameters resulted in seven parameters with heritabilities larger than 0.3 (Figure 3, right panel). We found that *BBslope* showed both low heritability and low CVs across all estimation cases. Despite exhibiting low variability across genotypes, *MRd* displayed relatively high heritability of 0.52. CO_2_ affinity for RuBisCO (*K_m_*-RuBisCO-CO_2_) had moderate heritability, which was notably higher than the heritability observed for the enzyme’s affinity to O_2_ (*K_m_*-RuBisCO-O_2_). Additionally, we found differences in the variability of V_max_ of RuBisCO carboxylation across genotypes between seasons, suggesting environment-specific influences on this key parameter.

In summary, increasing the number of jointly estimated parameters reduced uncertainty within individual genotypes but did not necessarily increase the parameter variability across genotypes. Furthermore, the highest mean heritabilities were observed when 11 parameters were jointly estimated. Together, these findings suggest that the estimated kinetic parameters are precise for a genotype, vary across genotypes, and are heritable, so they can be effectively used in downstream machine learning for GP.

### KineticGP outperforms classical GP for photosynthesis-related traits

In addition to the 68 genotypes, for which both *A*-*C_a_* and *A*-*PAR* curves were measured over two seasons, photosynthetic rates at saturating light were measured for 238 genotypes from the maize MAGIC population grown in 2022 and 2023 as well as 128 other genotypes grown in 2021. These genotypes were not included in the parameterization since no complete *A*-*C_a_* and *A*-*PAR* curves were available for 2022 and 2023. The photosynthesis rate at saturating light showed broad-sense heritability of 0.58 using all available data across three seasons. This indicates that the maximum predictability of photosynthetic rate at saturating light using genetic markers alone cannot exceed this value.

KineticGP relies on GP to build models for the *N* jointly estimated kinetic parameters using genetic markers from a training set of 68 genotypes. The models are then used to predict the *N* kinetic parameters for unseen genotypes (Figure 1d). The usage of the predicted kinetic parameters in the kinetic model in turn allows the simulation of photosynthetic rates under controlled gas exchange measurement conditions for unseen genotypes.

Before applying the approach to unseen genotypes, we assessed the predictability of kinetic parameters themselves through ten repetitions of 3-fold cross-validation using ridge regression best linear unbiased prediction (rrBLUP, Endelman, 2011). Among the predicted top 11 parameters, *τ_ActRubisco_* exhibited the highest median prediction accuracy (correlations of 0.21, Figure S15). In the case of 20 jointly estimated parameters, *MRd* and Perm-CO_2_ showed median accuracy above 0.15. These results were largely consistent with the estimated heritabilities of the parameters (Figure 3); namely, parameters with low heritability tended to show negative or near-zero predictive accuracy.

We then evaluated the ability of KineticGP to predict photosynthetic rate under saturating light for previously unseen genotypes under three prediction scenarios: (*i*) testing on seen 2022 season, (*ii*) testing on seen 2023 season, and (*iii*) testing on an unseen 2021 season (Figure S16). To this end, kinetic parameters for the unseen genotypes were predicted using the rrBLUP model, trained on the 68 genotypes with the jointly estimated parameters.

We compared two versions of KineticGP: KineticGP-11 and KineticGP-20, reflecting the number of jointly estimated kinetic parameters (*N*). All other kinetic parameters not included in the estimation retained their default values for predicting unseen genotypes. To enable comparisons with classical GP approaches, we also evaluated a baseline genomic prediction model, in which photosynthetic rate was predicted directly from genetic markers alone using rrBLUP (Figure S16).

Using the kinetic parameters predicted by rrBLUP, photosynthetic rate simulated at saturating light by KineticGP-11 achieved correlations of 0.22 and 0.18 with measured values for seasons 2022 and 2023, respectively (Figure 4a). In contrast, KineticGP-20 yielded near-zero correlations. The baseline models also performed poorly, yielding correlations of 0.074 and 0.16 using rrBLUP for seasons 2022 and 2023, respectively. Notably, none of the three approaches could predict photosynthetic rate for the 2021 season, underscoring the difficulty of generalizing across distinct environments. The latter finding also indicates that other environmental factors, now not included in the simulation, affect the transferability of the model to unseen seasons and contribute to the pronounced G×E interaction.

**Figure 4:**
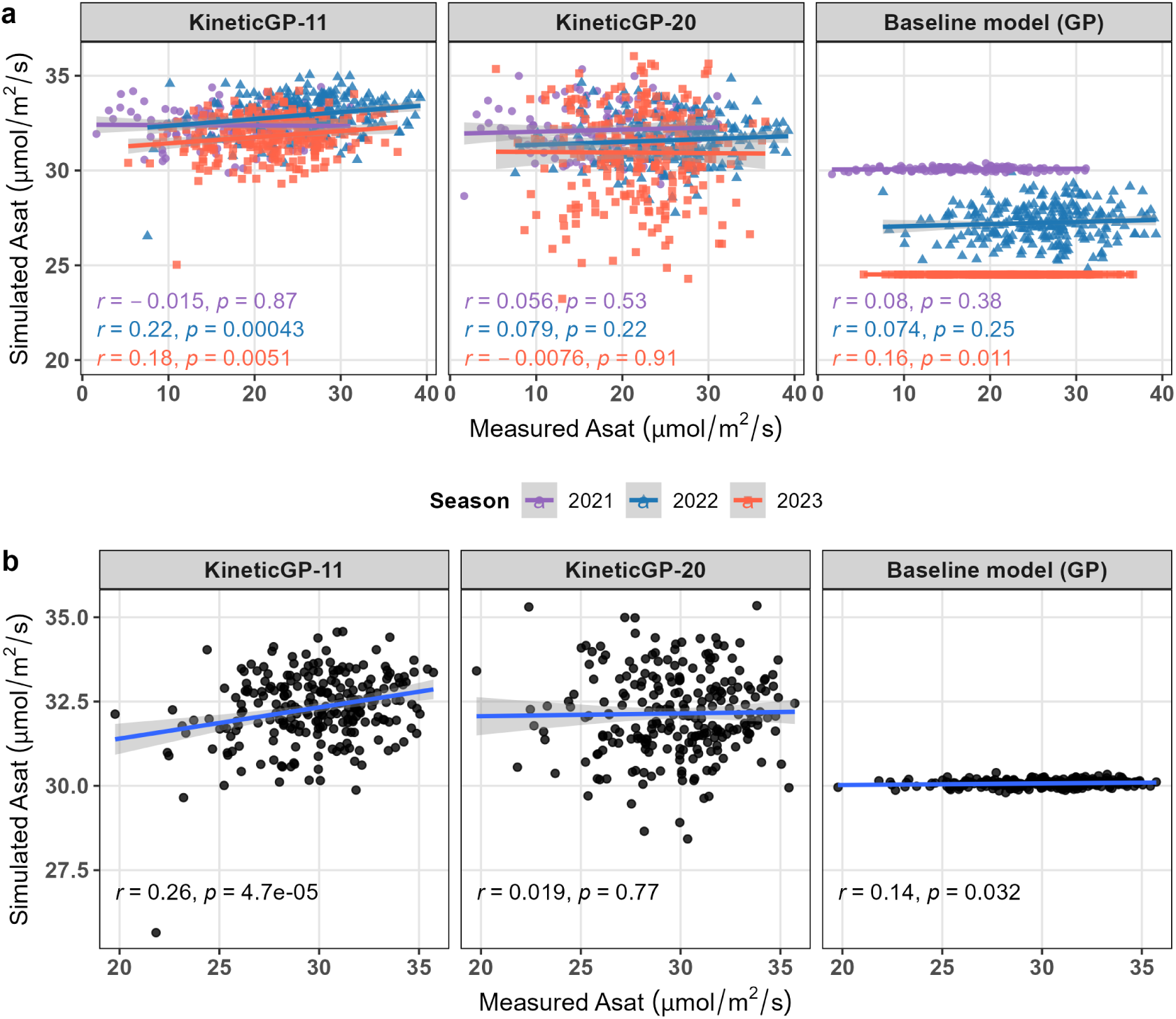
Comparison of performance for KineticGP versions and baseline GP model. (a) Simulated photosynthetic rates at saturating light (Asat) using KineticGP with 11 and 20 predicted parameters (KineticGP-11 and KineticGP-20) were compared against baseline model rrBLUP across individual seasons. The evaluation included 238 unseen genotypes grown in seen season (2022 and 2023) and 126 unseen genotypes from the unseen season (2021). Different colors represent the testing genotypes in the corresponding season. (b) Photosynthetic rates at saturating light (Asat) were simulated using KineticGP based on predicted BLUPs of season-specific V_max_ combined with other predicted parameters. For comparison, the baseline rrBLUP model was trained directly on the BLUPs of photosynthetic rate across the two seasons and applied for prediction. Both simulated and directly predicted rates were plotted against the BLUPs of measured Asat for seasons 2021 and 2023.

When rrBLUP models were trained to predict BLUPs of photosynthetic rates across two seasons, the resulting correlations for this baseline GP model improved moderately to 0.14 (Figure 4b). Incorporating G×E interactions in the Bayesian Generalized Linear Regression (BGLR, Pérez & de Los Campos, 2014) model led to a correlation of 0.17 with BLUPs across both seasons. To compare KineticGP in this setting, we first calculated the BLUP values of the season-specific *V_max_* parameters. These, along with other kinetic parameters, were then predicted using rrBLUP and used to simulate photosynthetic rates for unseen genotypes, as in the procedure above. Using this approach, KineticGP-11 yielded correlations of 0.26 with the BLUP of measured photosynthetic rates (Figure 4b).

Altogether, these results demonstrated the essential contributions of genotype-specific parameters for improving prediction accuracy of photosynthetic performance. Compared to baseline statistical models, even those accounting for G×E effects, KineticGP consistently yielded higher prediction accuracy for the 2022 and 2023 seasons by leveraging biologically grounded parameters and genotype-informed simulations that account for the effects of specific CO_2_ concentration and light intensity. However, consideration of other environmental factors, that affect photosynthesis-related traits, is warranted to improve predictions of KineticGP for the most challenging scenario with unseen genotypes in unseen environments.

## Conclusion

Here we introduced KineticGP, an innovative framework that integrates genotype-specific kinetic models with GP to simulate photosynthesis performance for genotypes and/or environmental conditions not included in the training set. The training population consisted of 68 genotypes from a maize MAGIC population, with gas exchange measurements obtained from varying levels of CO_2_ and light. These measured canonical curves allowed us to estimate genotype-specific kinetic parameters in kinetic model of C_4_ photosynthesis. The genotype-specific values were in turn used to train models for prediction of kinetic parameters. Therefore, KineticGP leverages the advantages of a mechanistic dynamic model, which, when combined with kinetic parameters predicted from genetic markers, improves the accuracy of predictions over classical GP for unseen genotypes in seen environments. Furthermore, KineticGP stands in contrast to the classical GP that is solely based on a machine learning model using genetic markers that results in environment-invariant predictions of phenotypic traits.

The application of KineticGP capitalized on our improvement of the default kinetic model of C_4_ photosynthesis (Wang et al., 2021), resulting from the update of the equilibrium constants using the eQuilibrator tool (Beber et al., 2022). The estimated genotype-specific kinetic parameters exhibited narrow confidence intervals for each genotype, indicating their precision and reliability for usage in downstream machine learning. Furthermore, we identified moderate variability across genotypes and moderate heritability for some of the kinetic parameters. These characteristics of the estimated parameters facilitated downstream predictions using SNP data. In comparison to the recently proposed C4TUNE (Wendering et al., 2025), an AI-driven approach for parameter prediction, the parameter estimation used in KineticGP imposes constraints on parameters that are not expected to change across seasons and has the ability of facile control on the number of parameters to be estimated, using the findings from sensitivity analyses. Future work will be aimed at fusing the ideas from KineticGP and C4TUNE.

To assess the predictive power of KineticGP for photosynthetic rates under saturating light, we evaluated its performance on unseen genotypes from both seen (2022 and 2023) and unseen (2021) growing seasons. Among the different KineticGP variants, the version estimating and predicting the top 11 kinetic parameters, selected based on the highest control coefficients, yielded the most accurate predictions for unseen genotypes from seen seasons. This finding suggests that measurements of few enzyme-specific parameters already suffices to improve predictions of photosynthesis-related traits. Our results demonstrated that KineticGP-11 significantly outperformed baseline GP models (using rrBLUP) across both seen seasons: (*e.g.*) increasing the predictability from 0.074 and 0.16 up to 0.22 and 0.18. Furthermore, when training was based on BLUPs of estimated *V_max_* parameters, simulations using the predicted kinetic parameters achieved a correlation of 0.26 with the BLUPs of measured photosynthetic rates from seasons 2022 and 2023. However, none of the approaches yielded significant correlations when applied to the unseen 2021 season.

These results highlight the advantage of incorporating genotype-specific kinetic model into genomic prediction framework, as opposed to relying solely on statistical models or generalized species-level mechanistic models. At the same time, these results reveal the difficulty of transferring model performance to entirely unseen seasons. Notably, the measured light-saturated photosynthetic rates showed only moderate correlations between 2021 and 2022 (r = 0.45), and between 2021 and 2023 (r = 0.46), indicating substantial seasonal variability. This observation indicates that different growth conditions may lead to acclimation effects and substantial GxE for photosynthesis. Whilst KineticGP can party capture G×E interactions by combining genotype-specific parameters with environmental input, full explanation of GxE variability may be limited by two factors: First, BLUPs of V_max_ parameters, that in part capture environmental effects, are used in GP to predict values for unseen genotypes; hence, the predicted values for V_max_ for unseen genotypes may not capture the season-specific protein allocation. To incorporate field conditions into the parameter estimation process, one must account for season-specific modulation of enzyme allocation, V_max_. Second, the current implementation of KineticGP uses the same initial metabolite concentrations across all genotypes at the first point of the simulated response curves (see Methods), which in part may affect the estimated parameter values. Nevertheless, we found that simulations from the second point of the response curves already start with different metabolite concentrations (Figure S17). Future work may consider acquiring empirical metabolite measurements that can be used in simulations.

In conclusion, KineticGP provides a template for hybrid GP that combines mechanistic and machine learning approaches. This framework can be employed to predict molecular and physiological traits related not only to metabolism, but also signalling and regulation, for which mechanistic models are actively developed.

## Materials and Methods

### Gas exchange measurements

The experimental data for this study were obtained from the field trials of the maize MAGIC population (Dell’Acqua et al., 2015) conducted at the National Institute of Agricultural Botany (NIAB, Cambridge, UK) over three consecutive seasons (2021, 2022, and 2023). The full details of the experimental design has previously been described in full detail (Ferguson et al., 2025). For this study, we made use of gas exchange measurements under varying ambient CO_2_ (*C_a_*) and photosynthetically active radiation (*PAR*), which can be represented as *A*-*C_a_* and *A*-*PAR* curves. *A*-*C_a_* curves were available for 78, 88 and 91 recombinant inbred lines with *A*-*C_a_* measurements in the three corresponding years, respectively. *A*-*PAR* measurements were only available for 314 genotypes from 2022 and 2023, while photosynthetic rate at saturating light were recorded for 151 lines in 2021. The estimation of kinetic parameters was conducted for 68 genotypes with both *A*-*C_a_* and *A*-*PAR* data from 2022 and 2023. The remaining data served as test sets to evaluate the performance of KineticGP.

The photosynthetic rate (*A*) and stomatal conductance (*g_s_*) were measured at twelve ambient CO_2_ concentrations (*C_a_*), starting at 400 µmol mol^-1^. Once *A* stabilized, measurements were recorded every 120 seconds as *C_a_* increased to 600, 800, 1000 and finally up to 1250 µmol mol^-1^. Afterward, *C_a_* was restored to 400 µmol mol^-1^, followed by 300, 250, 200, 100, 75 and finally 25 µmol mol^-1^. Throughout *A*-*C_a_* measurements, *PAR* was kept constant at 1800 µmol m^-2^ s^-1^ and the exchanger temperature was set to 25^◦^C. The *A*-*PAR* curves were measured under a constant ambient CO_2_ (400 µmol mol^-1^) and temperature (25^◦^C). The initial *PAR* level was set at 1800 µmol m^-2^ s^-1^ and then sequentially reduced to 1100, 500, 300, 150 and 50 µmol m^-2^ s^-1^.

### Kinetic model of C_4_-photosynthesis

The C_4_-photosynthesis model used in this study included mass balance of 109 metabolites whose concentration changes were determined by the 123 reactions in the system (Wang et al., 2021). The reaction rates involved 236 parameters, which we classified into four groups: (i) maximum velocities (V_max_), (ii) Michaelis-Menten constants (K_M_), (iii) activation rate constants of light-regulated enzymes, and (iv) the membrane permeability of specific metabolites. Since V_max_ values are the product of enzyme turnover number (k_cat_) and total enzyme concentration, this type of parameters can vary across different seasons. Thus, V_max_ parameters for 2022 and 2023 were treated as separate variables. All other parameters were required to be the same between the two years, in line with biophysical constraints. In addition, we observed discrepancies between the equilibrium constants used by Wang et al. (2021) and those provided by eQuilibrator (Beber et al., 2022). Therefore, we refined 29 out of the 35 equilibrium constants accordingly, while ensuring the stability of the dynamic system.

### Parameterization of genotype-specific kinetic models

The objective of model parameterization was to identify the set of kinetic parameters that results in simulated profiles (photosynthetic rate and stomatal conductance) as close as possible to the measured data. The standard distance metric used for fitting was the chi-square error (*χ*^2^), calculated using the formula:

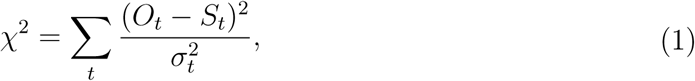

where *O_t_* and *σ_t_* denote the measured data and their standard deviation at a given time point, while *S_t_* is the simulation result at the same time point. The advantage of using this metric is that it allows us to determine if the resulting fit is statistically significant at a given significance level, defined by the degrees of freedom, which corresponds to the number of data points minus the number of fit parameters.

Optimization of the non-linear objective (*χ*^2^), which embeds ordinary differential equation (ODE) simulations and involves a large number of parameters, does not guarantee to reach the global optimum. This is a common challenge in optimization problems and various algorithms have been developed to address it. In this study, we used PESTO’s parallel tempering method for parameterization, and a simplified scheme of optimization is illustrated in Algorithm 1. PESTO is a Bayesian approach that provides a probability distribution for the fitted parameters allowing the most probable value to be selected (Liu et al., 2018). Like other Bayesian methods, PESTO also provides confidence intervals for sampled parameters, enhancing the reliability of the parameter estimates. The classical Markov-chain Monte Carlo (MCMC) method evaluates the system’s energy using a single stochastic process and accepts or rejects updates based on the temperature, which is an auxiliary variable of the sampling approach. At high temperature, the system explores larger space, while at low temperature, the system may become trapped in local energy minima. Parallel tempering (Geyer, 1991) was developed to address this issue by simulating replicas of the original system at different temperature and allowing exchange of complete configurations between systems, enabling systems at low temperature to escape local minima.

The required data for the optimization included:

1. Measured photosynthetic rates in response to environmental perturbations that reflect how a genotype responds to environmental changes:

a. mean and standard deviation of photosynthetic rate (**ACa**, *σ_ACa_*) and stomatal conductance (**g_s_**, *σ_g__s_*) across replicates at different CO_2_ concentrations(**C_a_**).
b. mean and standard deviation of photosynthetic rate (**APAR**,*σ_APAR_*) under varying photosynthetically active radiation levels (**PAR**).
2. Initial guess of kinetic parameters (**k_0_**), based on maize-specific parameters from Wang et al. (2021), which were obtained from literature references, adapted from other kinetic models, or estimated from model simulation.
3. Initial metabolite concentrations (**x_t0_**), required as an initial state of the ODE simulations. We used the initial metabolite concentrations that Wang et al. (2021) used for simulation. Further details on the simulation using these initial metabolite concentrations can be found in Supplementary Information A.2.

Simulation of photosynthetic rate and stomatal conductance was achieved by integrating the ODEs in the C_4_-photosynthesis model, given the initial metabolite concentrations (**x_t0_**), kinetic parameters (**k_sampled_**) and environmental factors (**C_a_** and **PAR**) as inputs. The simulation starts with *x_t_*_0_ and the first ambient CO_2_ level (**C_a_**(1)), allowing the system to reach steady state, at which CO_2_ assimilation and stomatal conductance were recorded as **ACa_sim_**(1) and **g_ssim_** (1), respectively. Subsequently, the ambient CO_2_ concentration was changed to **C_a_**(2) and the simulation was carried out for 120 seconds, to mimic observations, using the final metabolite concentrations from the first step. This process was repeated for the remaining measured **C_a_** levels, at constant mean chamber air temperature (*Tair*) for the given genotype. For simulation under changing *PAR*, a similar procedure was followed, with a constant air temperature of 25^◦^C. The simulated *A*-*C_a_*, *g_s_* and *A*-*PAR* curves were used to calculate the *χ*^2^ based on measured profiles. The photosynthesis simulation was integrated in the objective for MCMC sampling, as illustrated in the function ObjFunc in Algorithm 1.

The sampling algorithm started with the initial guess of kinetic parameters (**k_0_**) and calculated the *χ*^2^ between measured and simulated profiles. If a new set of sampled parameters resulted in a smaller *χ*^2^, they were included in the ensemble; otherwise, they could be accepted if a random probability is below acceptance probability prescribed by the system temperature and posterior value. The sampling and update of parameters were performed by parallel tempering algorithm. This process continued until a full ensemble of parameter sets was generated for each genotype.

For sensitivity analysis, each kinetic parameter was sampled independently using MCMC, generating 500 samples per genotype. The smallest *χ*^2^ was obtained (*fval_optimized_*) and the corresponding parameter value was selected (*x_optimized_*). The control coefficient was then calculated for each parameter as:

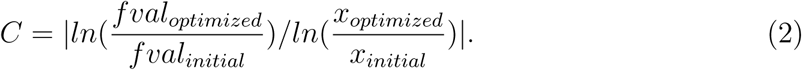

This coefficient quantifies the influence of each parameter on model fitting—higher values indicate a stronger impact.

The top *N* parameters with the highest control coefficients (N = 10, 20, 30 and 40) were selected for joint optimization. Including the V_max_ variable from both seasons led to final subsets of 11, 20, 34 and 44 parameters. These corresponded to optimization problems with 45, 36, 22 and 12 degrees of freedom, respectively.

Using the outlined MCMC approach, the set of kinetic parameters that produced a reduced chi-square value (*χ*^2^*/deg*(*N*)) closest to one was selected as the final parameter set for each genotype. The degrees of freedom (*deg*(*N*)) was defined as 56 data points (from *A* and *g_s_* at eleven *C_a_* levels and *A* at six *PAR* levels over two seasons) minus the number of estimated parameters (*N*). This set, denoted as (**SampledKs**_**G_j_**), defined the genotype-specific kinetic model that achieved the most statistically significant fit to the measured data. To assess estimation robustness, 300 additional parameter sets were sampled around the optimal solution to construct confidence intervals.

### Workflow of the KineticGP framework

Genomic data were obtained from Hobby et al. (2025) and included 70000 single-nucleotide polymorphisms (SNPs), encoded as −1, 0, and 1 for minor homozygous, heterozygous, and major homozygous at each loci, respectively. Narrow-sense heritability of the parameters was estimated using the heritability package in R (Kruijer et al., 2015). The required kinship matrix was generated from SNPs data using TASSEL5 (Bradbury et al., 2007). Broad-sense heritability of photosynthetic rate at saturating light, considering all genotypes across three seasons, was calculated using software package META-R (Alvarado et al., 2020).

To evaluate the average predictability of individual kinetic parameters, we performed ten iterations of 3-fold cross-validation using ridge regression best linear unbiased prediction (rrBLUP, Endelman, 2011), estimating marker effects from SNPs data. The prediction accuracy of the kinetic parameters was quantified by calculating the Pearson correlation coefficient between estimated and predicted values across each iteration.

The generalizability of KineticGP to unseen genotypes or environment was compared with baseline models, that predict photosynthetic rate directly from SNPs data using rrBLUP, across two testing scenarios.

1. Testing on seen seasons with previously unseen genotypes. The testing set comprises 238 genotypes with measured photosynthetic rates under light-saturating conditions in 2022 and 2023. The kinetic parameters for these testing genotypes were predicted using rrBLUP models trained on the 68 parameterized genotypes. The kinetic model then simulate photosynthetic rate using the predicted parameters. The baseline models predicted photosynthetic rate directly using SNPs for each season separately.
2. Testing on unseen season with previously unseen genotypes. Since *A*-*PAR* curves were not measured in season 2021, this season was excluded during estimation of kinetic parameters. There were 128 unseen genotypes with measured light-saturating photosynthetic rate in 2021. In this scenario, rrBLUP was trained using the Best Linear Unbiased Predictors (BLUPs) of estimated V_max_ parameters across the 2022 and 2023 seasons, along with the estimated season-invariant parameters as the response variables. Then the predicted parameters for the testing set were used to simulate photosynthetic rate for these 128 genotypes. For the baseline models, the response variable was the BLUPs of measured photosynthesis rates across both seasons.

Unlike the baseline models, KineticGP offers greater flexibility in terms of the usage of predicted kinetic parameters. The C_4_ photosynthesis model contains in total 236 kinetic parameters and through sensitivity analysis, top 10 and 20 kinetic parameters with the highest control coefficient were identified. Including the *V_max_* variables from both seasons resulted in final parameter sets of 11 and 20 parameters, respectively. These were used for estimation and downstream genomic prediction, and are referred to as KineticGP-11 and KineticGP-20. The simulation using predicted parameters was performed under gas exchange measurement condition, in which temperature and CO_2_ levels were kept constant for all genotypes, at saturating-light level.

## Author contribution

R.X. contributed methodology, software, investigation, visualization, writing. J.F. contributed investigation, data acquisition and curation. D.H. contributed software, investigation. M.R. contributed software, investigation. P.W. contributed software, investigation. J.K. contributed conceptualization, writing, supervision, project administration, funding acquisition. Z.N. contributed conceptualization, methodology, writing, supervision, project administration, funding acquisition.

## Data and code availability

The gas exchange measurements for maize genotypes are available at Zenodo repository (https://doi.org/10.5281/zenodo.15966533). Part of these data has been used to estimate photosynthesis-related traits, that are in turned modeled using hyperspectral reflectance data in another study (Xu et al., 2025). All codes and data to ensure reproducibility of the results can be accessed at: https://github.com/Rudan-X/KineticGP

## A. Supplementary Information

### A.1 Update of equilibrium constants using eQuilibrator database

Since the equilibrium constants (K_eq_) are determined by thermodynamic principles, they are not genotype-specific and therefore were excluded from the estimation and prediction. The kinetic model of Wang et al. (2021) contains 35 equilibrium constants which were collected from 15 publications (Colowick & Sutherland, 1942; Copper & Meister, 1972; Datta & Racker, 1961; Espada, 1962; Flodgaard & Fleron, 1974; Guynn, 1982; Hansen et al., 1966; Harary et al., 1953; Keirns & Wang, 1972; Kleczkowski et al., 1985; Knaff, 1996; Laisk & Edwards, 2000; Lunn & ap Rees, 1990; Pocker & Miksch, 1978). These studies derived the equilibrium constant by calculating the Gibbs energy based on measured concentrations during enzymatic essays, which used proteins extracted from different species (yeast, plant tissues, and animal organs). In order to obtain a more consistent set of equilibrium constants, we extracted the equilibrium constants from the eQuilibrator database (Beber et al. (2022), version 3.0, K_eq_ adjusted to Mg, ionic strength and pH), which has been shown to provide accurate estimates of standard Gibbs free energy.

The updated kinetic model consistently yielded lower *χ*^2^ errors compared to that using *K_E_* values provided by Wang et al. (2021), with significant improvements observed for *A*-*C_a_* and *A*-*PAR* curves in both seasons (Figure S17). For instance, updated models led to median *χ*^2^ values of 31.27 and 42.14 for *A*-*C_a_* curves in seasons 2022 and 2023, respectively, compared to 42.62 and 63.86 with the original models. Notably, the *χ*^2^ error from *A*-*C_a_* curves were higher than for *A*-*PAR* curves, while the stomatal conductance profiles showed the largest discrepancy between initial simulations and measurements. These findings show that genotype-specific kinetic values are needed to improve the fit of measured data and simulated photosynthesis-related traits.

### A.2 The usage of initial metabolite concentrations

Since we lacked access to the metabolic state (*i.e.*, metabolite concentrations) of different genotypes across environments, we initialized the model simulation using the same metabolite concentrations for all genotypes at the first measured point. For *A*-*C_a_* curves, the model was first simulated atr *C_a_* = 400 µmol CO_2_ mol^-1^, using the initial concentrations from Wang et al. (2021), until *A* reaches a steady-state. The resulting metabolite concentrations were then used as initial states for simulating the next *C_a_* level (600 µmol CO_2_ mol^-1^). The same approach was applied for simulating *A − PAR* curves. To demonstrate that the metabolic states used as initial conditions at *C_a_* of 600 µmol CO_2_ mol^-1^ vary between genotypes, we determined the variation in the concentration of the metabolites in the model over the different genotypes (Figure S18). We found that there is a considerable variability (one order of magnitude difference) between the initial concentrations of selected metabolites at *C_a_* level of 600 µmol CO_2_ mol^-1^, demonstrating that we can rely on the modification of parameters to achieve reliable simulations.

## Supplementary Tables

**Table S1:** List of reactions and their corresponding kinetic parameters. The top 44 kinetic parameters with the highest control coefficients are highlighted in bold.

| Reactions | Enzymes | $K_m, K_i$ | $V_{max}$ |
| --- | --- | --- | --- |
| 1. CO<sub>2</sub>->HCO<sub>3</sub> [MC] | 4.2.1.1 | **Km1-CO2** | Vm1 |
| 2. HCO<sub>3</sub>->PEP+OAA [MC] | 4.1.1.31 | **Km2-HCO3, Km2-PEP**<br>Ki2-MAL,Ki2-MALn | **Vm2** |
| 3. OAA+NADPH->MAL+NADP [MCchl] | 1.1.1.82 | Km3-NADPH,Km3-OAA<br>Km3-NADP,Km3-MAL | **Vm3** |
| 4. MAL+NADP->CO<sub>2</sub>+PYR+NADPH [BCchl] | 1.1.1.40 | Km4-CO<sub>2</sub>,Km4-NADP<br>Km4-NADPH,Km4-Pyr<br>Km4-MAL | **Vm4** |
| 5. PYR+ATP->PEP [MCchl] | 2.7.9.1 | Ki5-PEP,Km5-ATP<br>Km5-Pyr | **Vm5** |
| 6. CO<sub>2</sub>+RuBP->2PGA [BCchl] | 4.1.1.39 | **Km6-CO2,Km6-O2**<br>**Km6-RuBP**,Ki6-PGA<br>Ki6-FBP,Ki6-SBP<br>Ki6-Pi,Ki6-NADPH | **Vm6** |
| 7. PGA+ATP->ADP+DPGA [BCchl] | 2.7.2.3 | Km7-ATP, **Km7-PGA** | **Vm7-8** |
| 8. DPGA+NADPH->GAP+Pi+NADP [BCchl] | 1.2.1.13 | Km8-NADPH |  |
| 9. GAP<->DHAP [BCchl] | 5.3.1.1 |  |  |
| 10. DHAP+GAP<->FBP [BCchl] | 4.1.2.13 | **Km10-DHAP**,Km10-FBP<br>**Km10-GAP** | Vm10 |
| 11. FBP<->F6P+Pi [BCchl] | 3.1.3.11 | Ki11-F6P,Ki11-Pi<br>Km11-FBP | Vm11 |
| 12. E4P+DHAP<->SBP [BCchl] | 4.1.2.13 | **Km12-DHAP,Km12-E4P** | **Vm12** |
| 13. SBP<->S7P+Pi [BCchl] | 3.1.3.37 | Ki13-Pi,Km13-SBP | Vm13 |
| 14. F6P+GAP<->E4P+Xu5P [BCchl] | 2.2.1.1 | Km14-E4P,Km14-F6P<br>**Km14-GAP**,Km14-Xu5P | Vm14 |
| 15. S7P+GAP<->Ri5P+Xu5P [BCchl] | 2.2.1.1 | **Km15-GAP**,Km15-Ri5P<br>Km15-S7P,Km15-Xu5P | Vm15 |
| 16. Ri5P<->Ru5P [BCchl] | 5.3.1.6 |  |  |
| 17. Xu5P<->Ru5P [BCchl] | 5.1.3.1 |  |  |
| 18. Ru5P+ATP<->RuBP+ADP [BCchl] | 2.7.1.19 | **Km18-ATP,Km18-Ru5P**<br>Ki18-ADP,Ki18-ADP2<br>Ki18-PGA, **Ki18-Pi**<br>Ki18-RuBP | Vm18 |
| 19. PGA+ATP->ADP+DPGA [MCchl] | 2.7.2.3 | Km19-ATP,Km19-PGA | Vm19-20 |
| 20. DPGA+NADPH->GAP+Pi+NADP [MCchl] | 1.2.1.13 | Km20-NADPH |  |
| 21. F6P<->G6P, G6P<->G1P [BC] | 5.3.1.9<br>5.4.2.2 |  |  |
| 22. PGA->sink | PGA<br>sink | Km22-PGA | Vm22 |
| 23. DHAP+GAP<->FBP [MC] | 4.1.2.13 | Km23-DHAP,Km23-GAP<br>Km23-FBP | Vm23 |
| 24. FBP<->F6P+Pi [MC] | 3.1.3.11 | Ki24-F26BP,Ki24-F6P<br>Ki24-Pi,Km24-FBP | Vm24 |
| 25. F6P<->G6P, G6P<->G1P [MC] | 5.3.1.9<br>5.4.2.2 | Ki24-F26BP,Ki24-F6P<br>Ki24-Pi,Km24-FBP |  |
| 26. G1P+UTP<->UDPG+PPi [MC] | 2.7.7.9 | Km26-G1P,Km26-PPi<br>Km26-UDPG,Km26-UTP | Vm26 |

| Reactions | Enzymes | K<sub>m</sub>, K<sub>i</sub> | V<sub>max</sub> |
| --- | --- | --- | --- |
| 27. UDPG+F6P<->SUCP+UDP [MC] | 2.4.1.14 | Km27-F6P,Km27-UDPG<br>Ki27-FBP,Ki27-Pi<br>Ki27-Suc,Ki27-SucP<br>Ki27-UDP | Vm27 |
| 28. SUCP<->Pi+SUC [MC] | 3.1.3.24 | Km28-Suc,Km28-SucP | Vm28 |
| 29. SUC->sink [MC] | Sucrose<br>sink | Km29-Suc | Vm29 |
| 30. F6P+ATP->F26BP+ADP [MC] | 2.7.1.105 | Km30-ATP,Km30-F26BP<br>Km30-F6P ,Ki30-ADP<br>Ki30-DHAP | Vm30 |
| 31. F26BP[c]->F6P[c]+Pi[c] [MC] | 3.1.3.46 | Ki31-F6P,Ki31-Pi<br>Km31-F26BP | Vm31 |
| 32. ADP+Pi<->ATP [MCthy] | 3.6.3.14 | Km32-ADP,Km32-ATP<br>Km32-PiX32,Y32<br>F32,Q32,D32 | **J<sub>max</sub>**<br>Vm32 |
| 33. NADP<->NADPH [MCthy] | 1.18.1.2 | **Km33-NADP**<br>E33,Km33-NADPH | **Vm33** |
| 34. ADP+Pi<->ATP [BCthy] | 3.6.3.14 | Km34-ADP,Km34-Pi<br>Km34-ATP,G34 | Vm34 |
| 35. Metabolite transport<br>through plasmodesmata |  | Km35Mc-OAA<br>Ki35Mc-mal-OAA<br>Km35M-mal<br>Ki35M-OAA-mal<br>Km35B-mal,Km35B-pyr<br>Km35M-pyr,Km35M-PEP | **Vm35Mc-OAA**<br>Vm35B-pyr<br>Vm35M-pyr<br>Vm35M-PEP<br>Vm35B-PEP<br>Vm35B-mal<br>Vm35C-mal<br>Vm35-Hep |
| 36. PPi + H2O <-> 2 Pi | 3.6.1.1 |  |  |
| 37. NADP<->NADPH [BCthy] | 1.18.1.2 | Km37-NADP<br>Km37-NADPH | Vm37 |
| 38. O2+RuBP<->PGCA+PGA [BCchl] | 4.1.1.39 | Km6-CO2,Km6-O2<br>Km6-RuBP,Ki6-PGA<br>Ki6-FBP,Ki6-SBP<br>Ki6-Pi,Ki6-NADPH | Vm38 |
| 39. PGCA->Pi+GCA [BCchl] | 3.1.3.18 | Km39-PGCA,Ki39-PI<br>Ki39-GCA | Vm39 |
| 40. GCA+O2<->H2O2+GOA [BCper] | 1.1.3.15 | KmGCA40 | Vm40 |
| 41. GOA+GLU<->GLY+KG [BCper] | 2.6.1.4 | Km41-GOA,Km41-GLU<br>Ki41-GLY | Vm41 |
| 42. GLY+NAD<->SER+NADH+NH3 | 1.4.4.2<br>2.1.2.1 | Km42-GLY,Ki42-SER | Vm42 |
| 43. SER+GOA<->HPR+GLY [BCper] | 2.6.1.45 | Km43-GOA<br>Km43-SER<br>Km43-GLY | Vm43 |
| 44. HPR+NAD<->GCEA+NADH [BCper] | 1.1.1.29 | Ki44-HPR,Km44-HPR | Vm44 |
| 45. GCEA+ATP<->PGA+ADP [BCper] | 2.7.1.31 | Km45-ATP,Km45-GCEA<br>Ki45-PGA | Vm45 |
| 46. GCA[BCper]<->GCA [BCchl] | GCA<br>transport | Km46-GCA,Ki46-GCEA | Vm46 |

| Reactions | Enzymes | $K_m$, $K_i$ | $V_{max}$ |
| --- | --- | --- | --- |
| 47. GCEA[BCper]<->GCEA [BCchl] | GCEA transport | Km47-GCEA,Ki47-GCA | Vm47 |
| 48. PGA<->PEP [MC] | 5.4.2.1<br>4.2.1.11 | Km48-PGA,Km48-PEP | Vm48 |
| 49. PPDK inactivation [MCchl] | PPDK<br>Regulatory<br>Protein | Kcat49-Inact,Km49-ADP<br>Ki49-Pyr,Km49-E |  |
| 50. PPDK activation [MCchl] | PPDK<br>Regulatory<br>Protein | Kcat50-Act,Km50-Pi<br>Ki50-AMP,Km50-EP<br>Ki50-ADP,Ki50-PPI |  |
| 51. G1P + ATP <-> PPI + ADPG [Bchl] | 2.7.7.27 | Ka51-PGA,Km51-G1P<br>,Km51-ATP,Ki51-APi-ATP<br>Km51-PPI,Ki51-CPP1-ATP<br>Km51-ADPG<br>Ki51-ADP-ATP | Vm51 |
| 52. PPI + H2O <-> 2 Pi [Bchl] | 3.6.1.1 | Km52-PPI | Vm52 |
| 53. ADPG <-> Starch [Bchl] | 2.4.1.21 | Km53-ADPG | Vm53 |
| 54. Hexose phosphate pool [Bchl] |  | Km54-pi,Km54-hexp |  |
| 55. PGA, GAP and DHAP transport [BC] | Triose<br>phosphate<br>translocator | Km55-PGA,Km55-GAP<br>Km55-DHAP | Vm55 |
| 56. PGA, GAP and DHAP transport [MC] | Triose<br>phosphate<br>translocator | Km56-PGA,Km56-GAP<br>Km56-DHAP | Vm56 |
| 57. Gs dynamics [MC] | Dynamic<br>stomatal<br>conductance | **Ki57**,Kd57<br>**BBslope**, BBintercept |  |
| Light-dependent enzyme activation | | **KaRac** | τPEPC<br>τFBPase<br>τSBPase<br>τATPsynthase<br>τGAPDH<br>τPRK<br>τRca<br>τNADPMDH<br>τ**RuBisCO** |
| Metabolite permeability |  | Perm-MAL<br>Perm-PYR<br>**PermMC-CO2**<br>Perm-PGA<br>PermBC-CO2<br>**gm** |  |

### Algorithm 1 Estimation of *N* kinetic parameters from C_4_ model for a given genotype *G_j_*

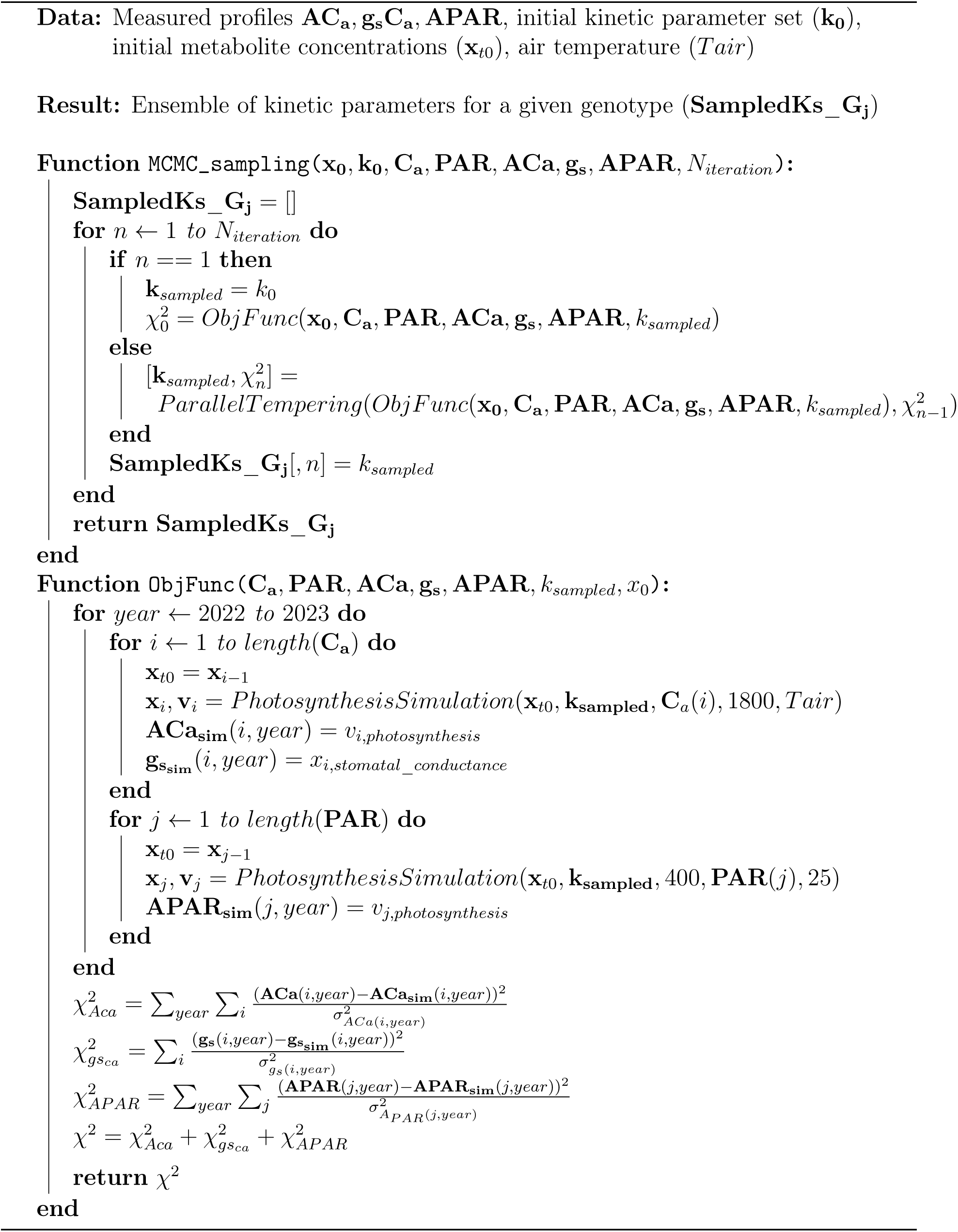

## Supplementary Figures

**Figure S1:**
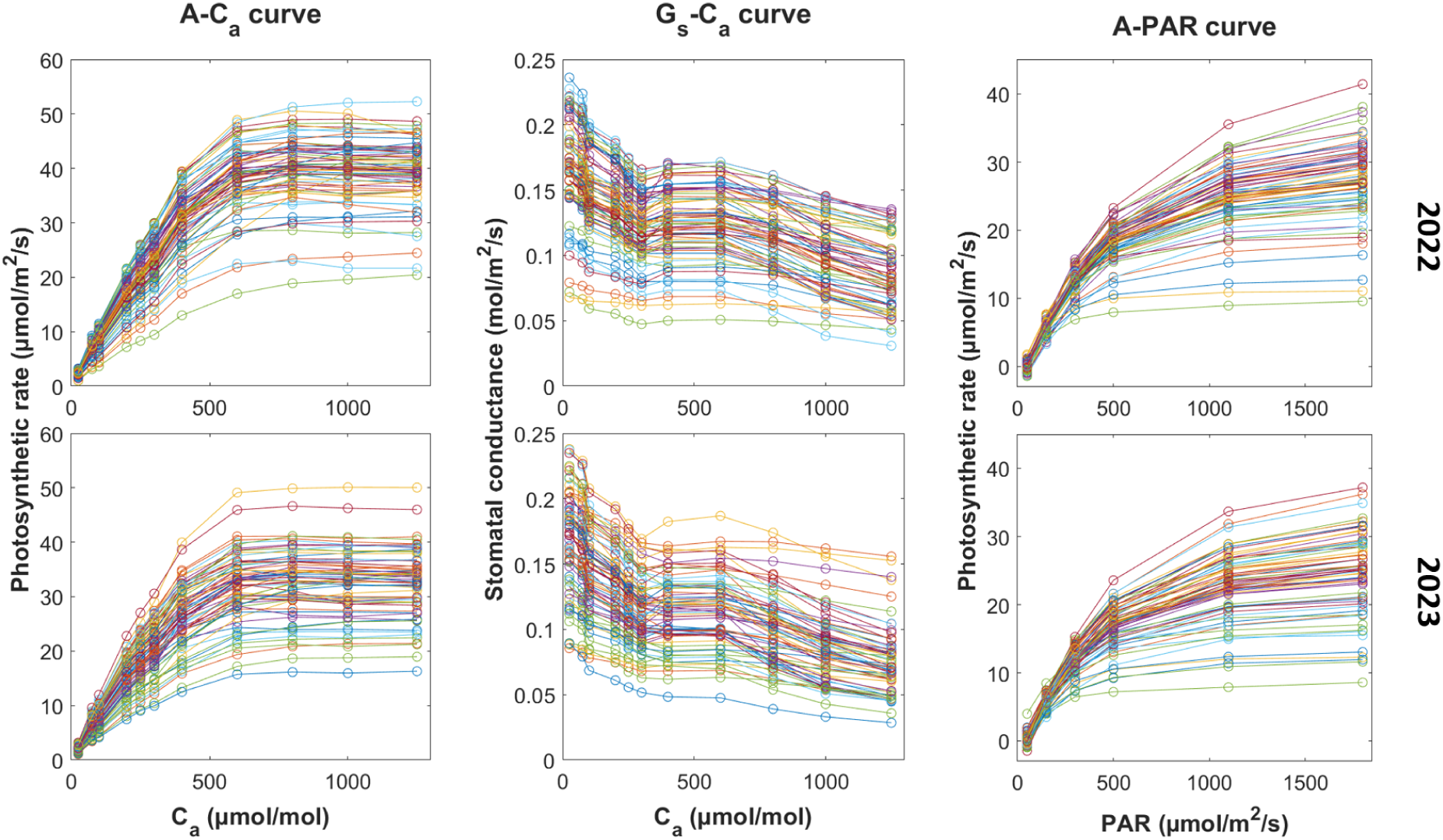
Characteristics of measured data sets. Measured photosynthetic rate (*A*) and stomatal conductance (*g_s_*) under changing ambient CO_2_ (*C_a_*) and photosynthetically active radiation (*PAR*) levels for 68 MAGIC maize recombinant inbred lines, grown in seasons 2022 and 2023.

**Figure S2:**
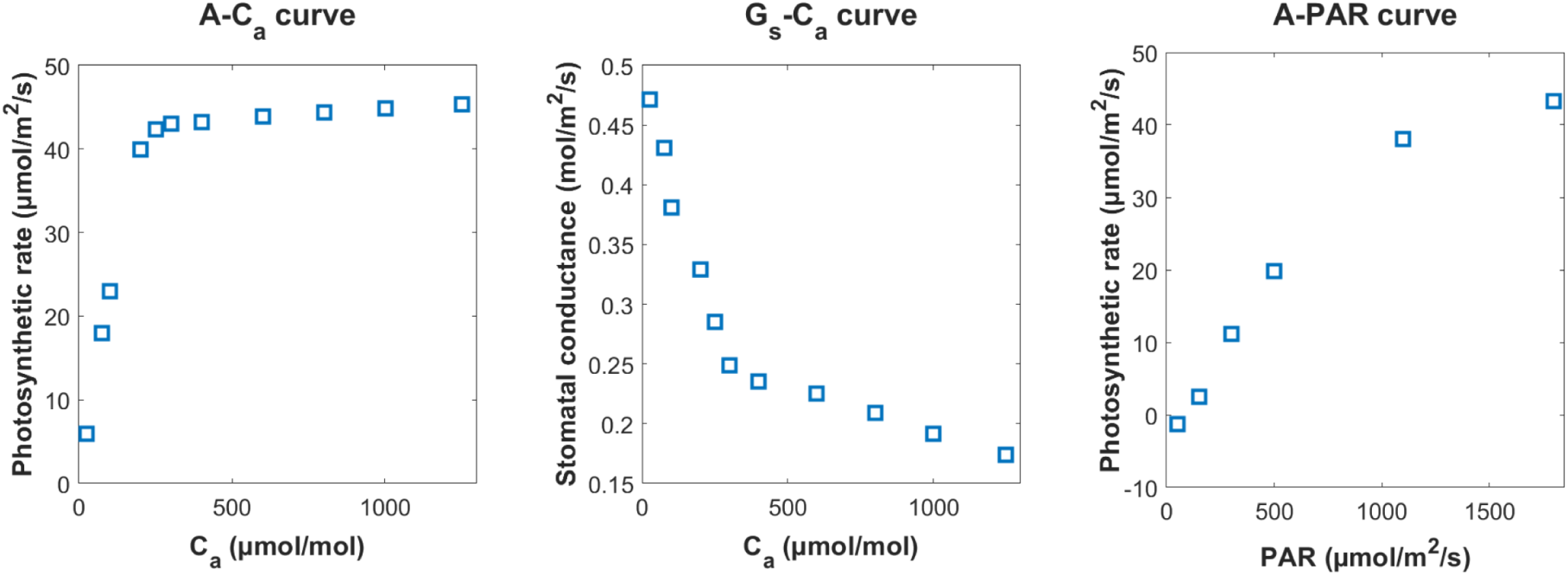
Simulated photosynthetic rate and stomatal conductance under changing ambient CO_2_ and PAR levels using maize-specific kinetic parameters from Wang et al. (2021).

**Figure S3:**
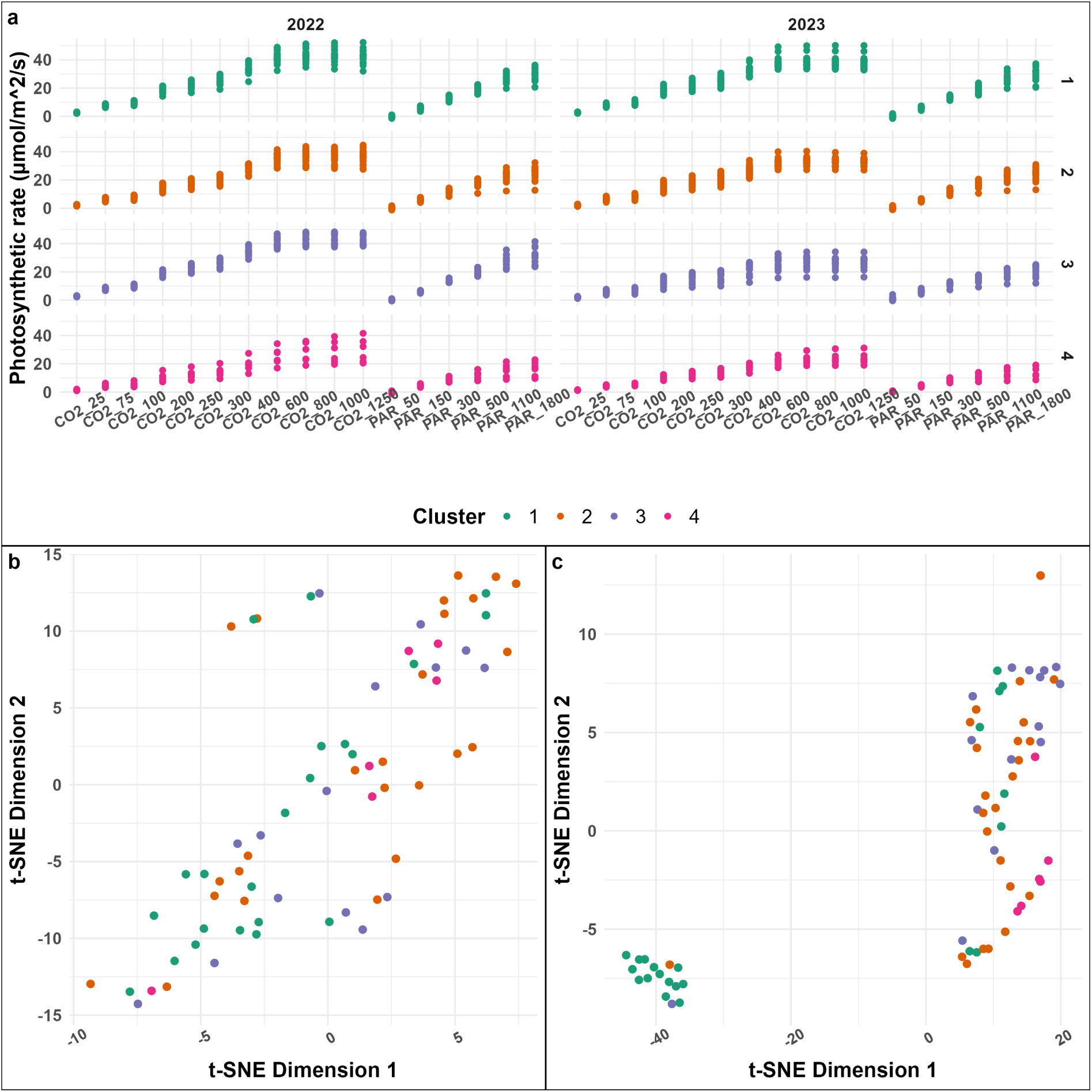
Clustering of 68 genotypes according to photosynthetic profile and kinetic parameters. (a) k-means clustering of genotypes in terms of their photosynthetic response to CO_2_ and PAR levels across both seasons. (b) Projection of all kinetic parameters in two t-SNE dimensions, the colors represent the same k-means clustering as in (a). (c) Projection in two t-SNE dimensions of 40 kinetic parameters with the highest control coefficient.

**Figure S4:**
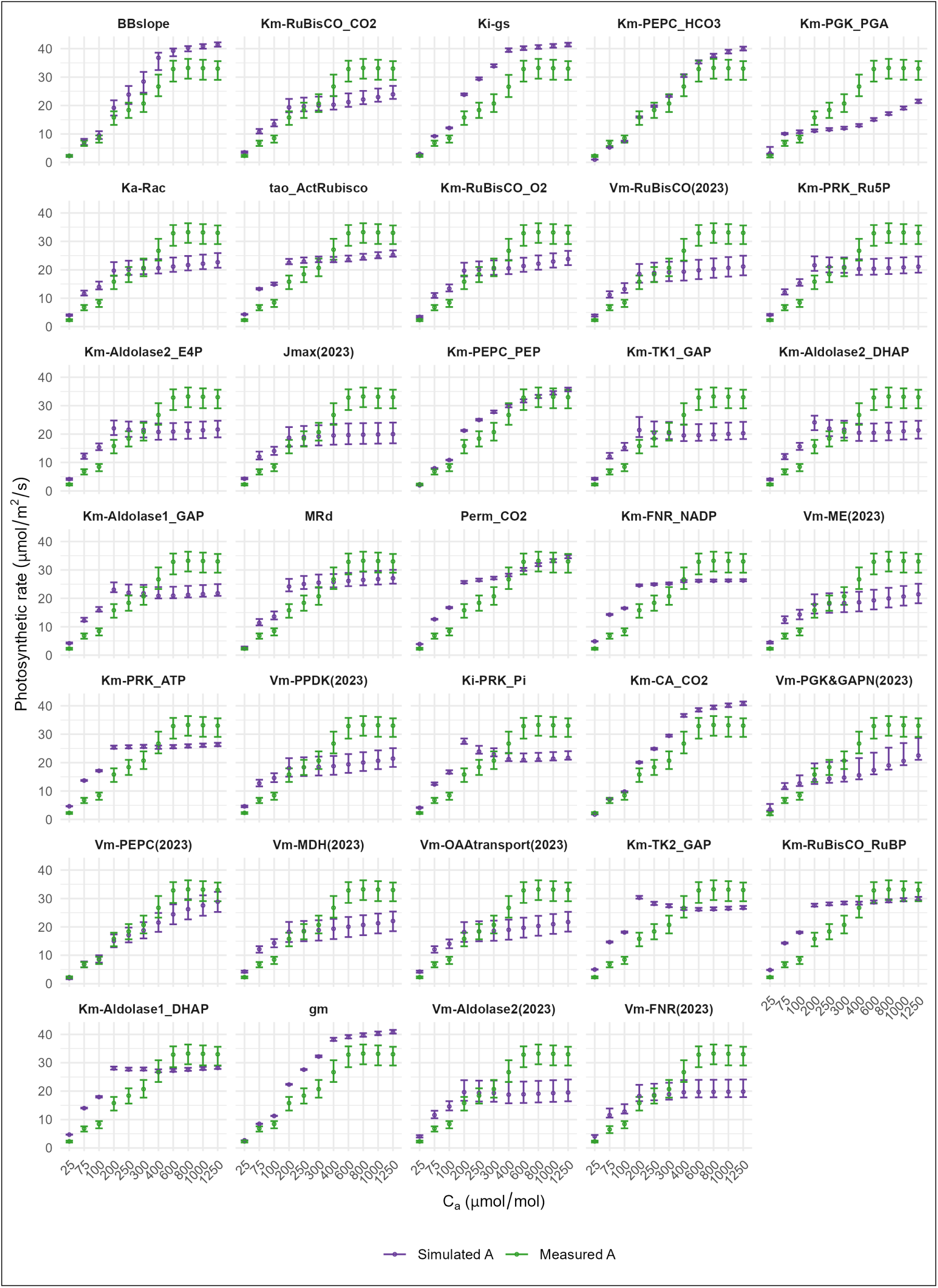
Comparison between simulated *A*-*C_a_* curves for season 2023 using individually estimated top-ranked kinetic parameters and measured *A*-*C_a_* curves. The median simulated *A* values across 68 genotypes (purple circles) at different *C_a_* levels were compared with the median measured *A* values (green circles). The error bars represent the inter-quartile range across genotypes, capturing the variability.

**Figure S5:**
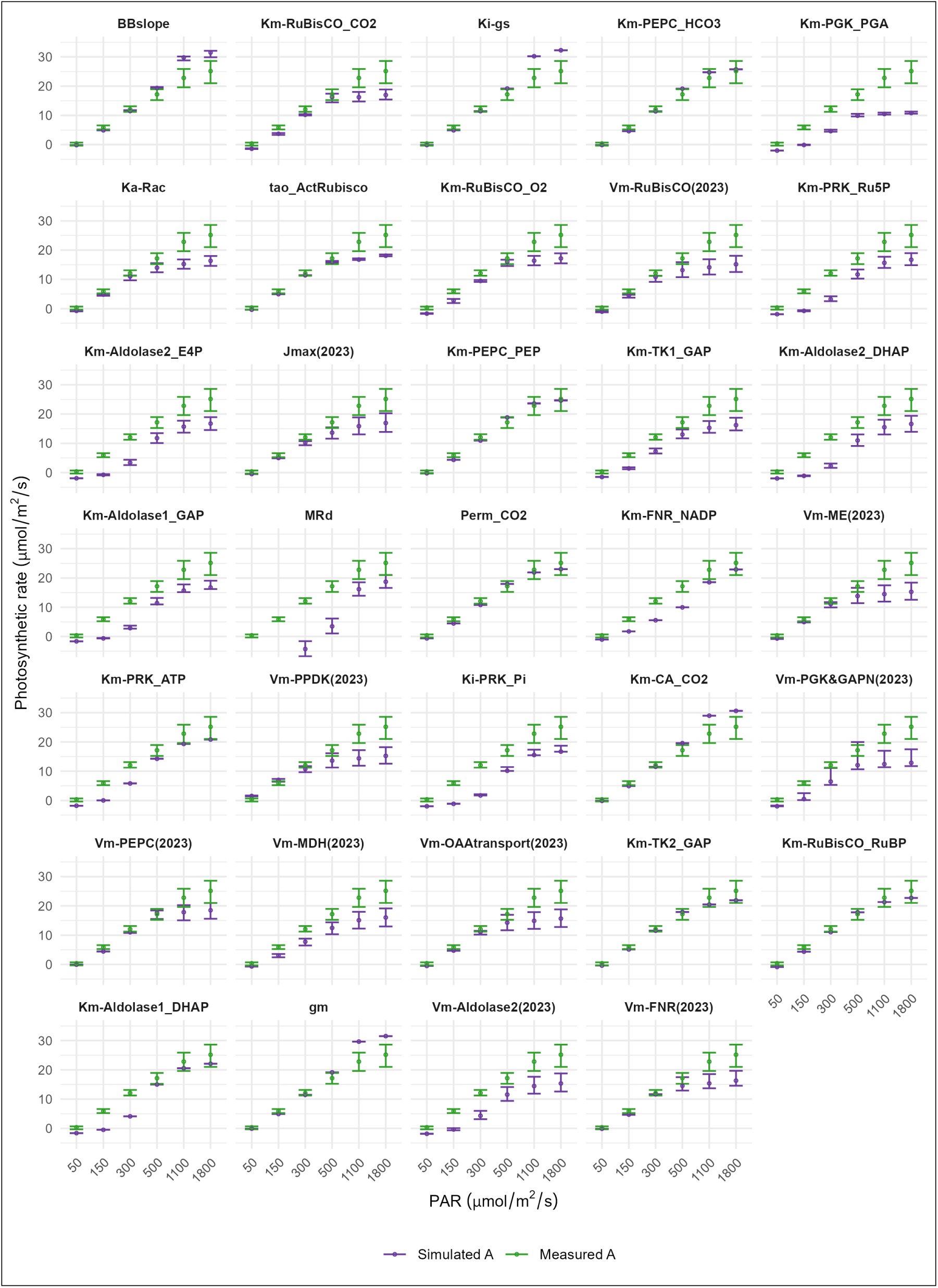
Comparison between simulated *A*-*PAR* curves for season 2023 using individually estimated top-ranked kinetic parameters and measured *A*-*PAR* curves. The median simulated *A* values across 68 genotypes (purple circles) at different *PAR* levels were compared with the median measured *A* values (green circles). The error bars represent the inter-quartile range across genotypes, capturing the variability.

**Figure S6:**
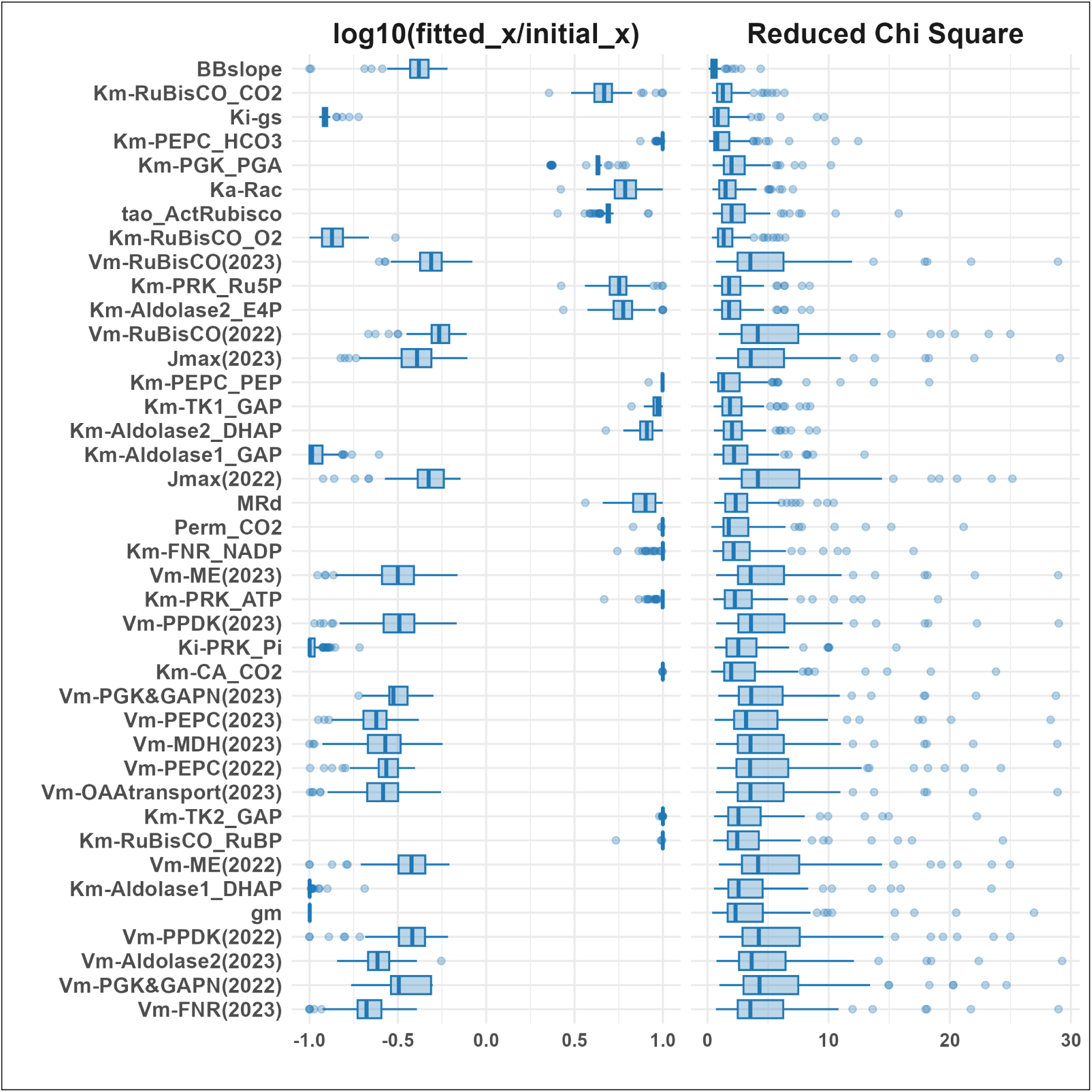
Distribution and fitting performance of kinetic parameters with the highest control coefficient. Left panel: logarithmic scale of the ratio between fitted and initial parameter values. Right panel: reduced *χ*^2^ values reflecting the fitting performance of each parameter based on individually optimized values. The boxplots represent the variability across 68 genotypes.

**Figure S7:**
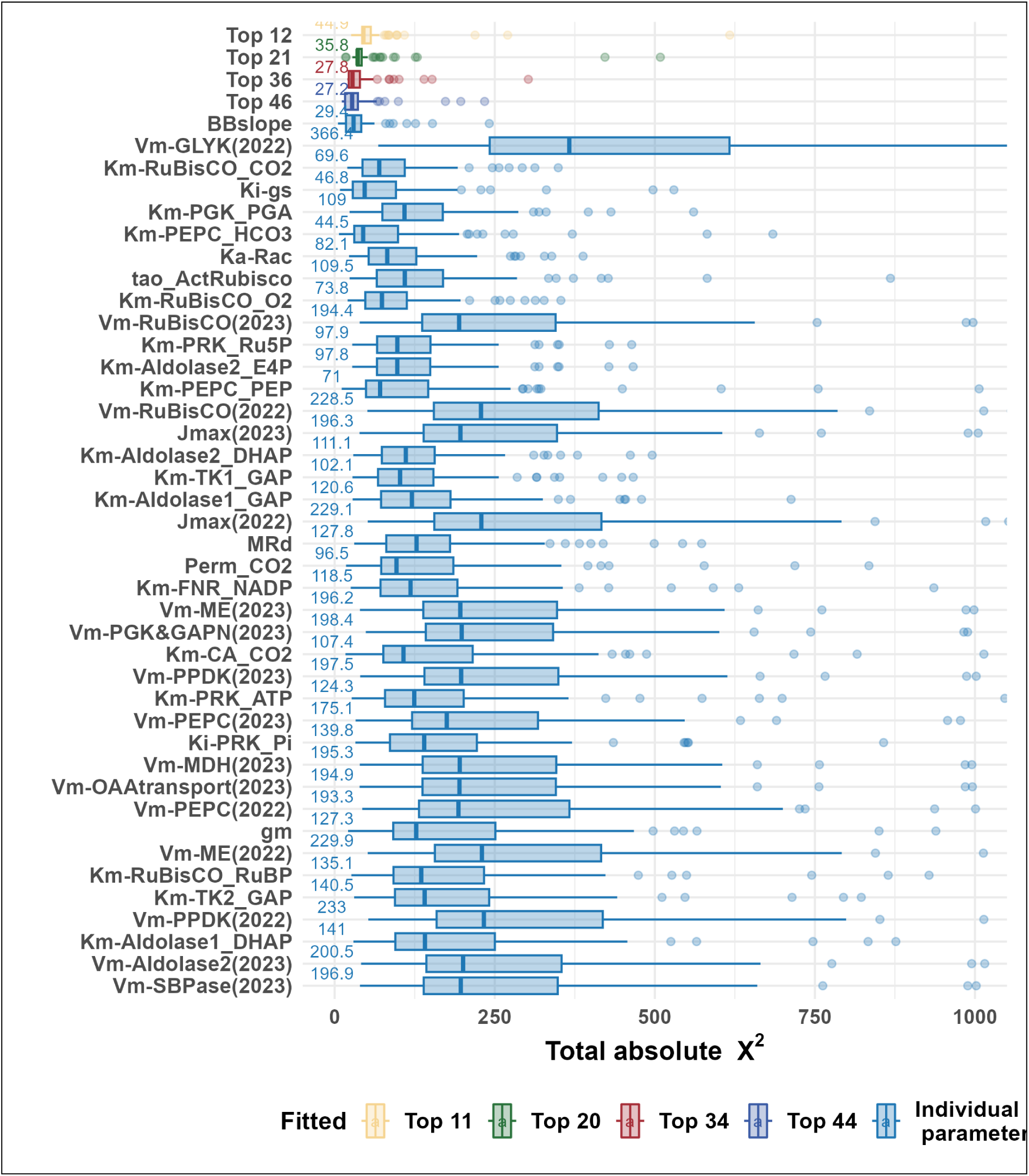
Comparison of total *χ*^2^ statistics of *A*-*C_a_*, *g_s_*-*C_a_* and *A*-*PAR* curves in two seasons between individually fitted parameters and selectively fitted parameters with the highest control coefficient.

**Figure S8:**
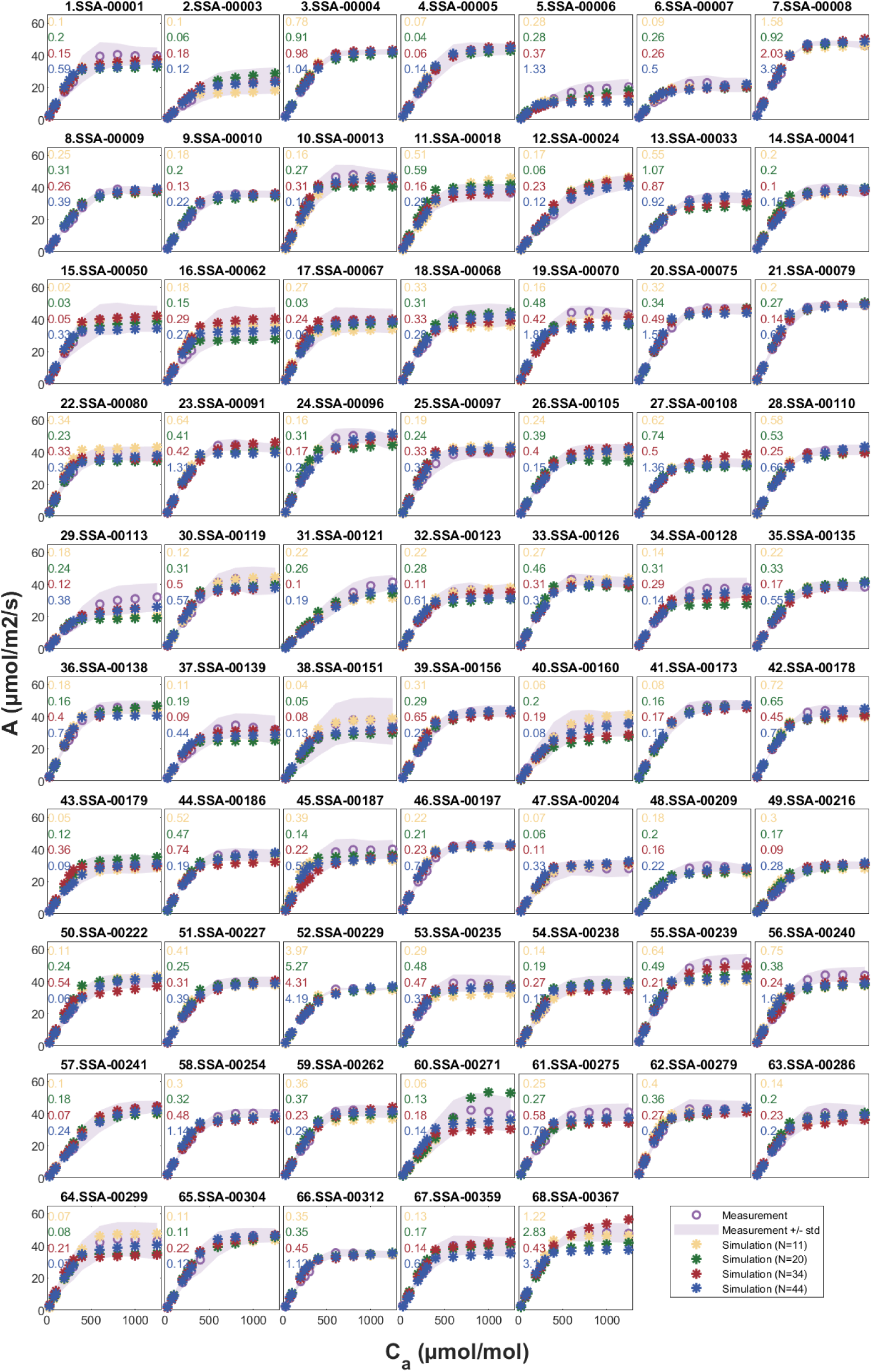
Fitting of photosynthetic rate (*A*) under varying ambient CO_2_ levels (*C_a_*) for 68 genotypes grown in 2022. Mean measured photosynthetic rate across replicates (circles) together with the standard deviation around (shaded area) are compared with the simulated values (stars), with different colors representing the number of estimated kinetic parameters: top 12 parameters (yellow), 21 parameters (green), 36 parameters (red) and 46 parameters (blue) with the highest control coefficient. The absolute *χ*^2^ statistics was annotated with color corresponding to different estimation cases.

**Figure S9:**
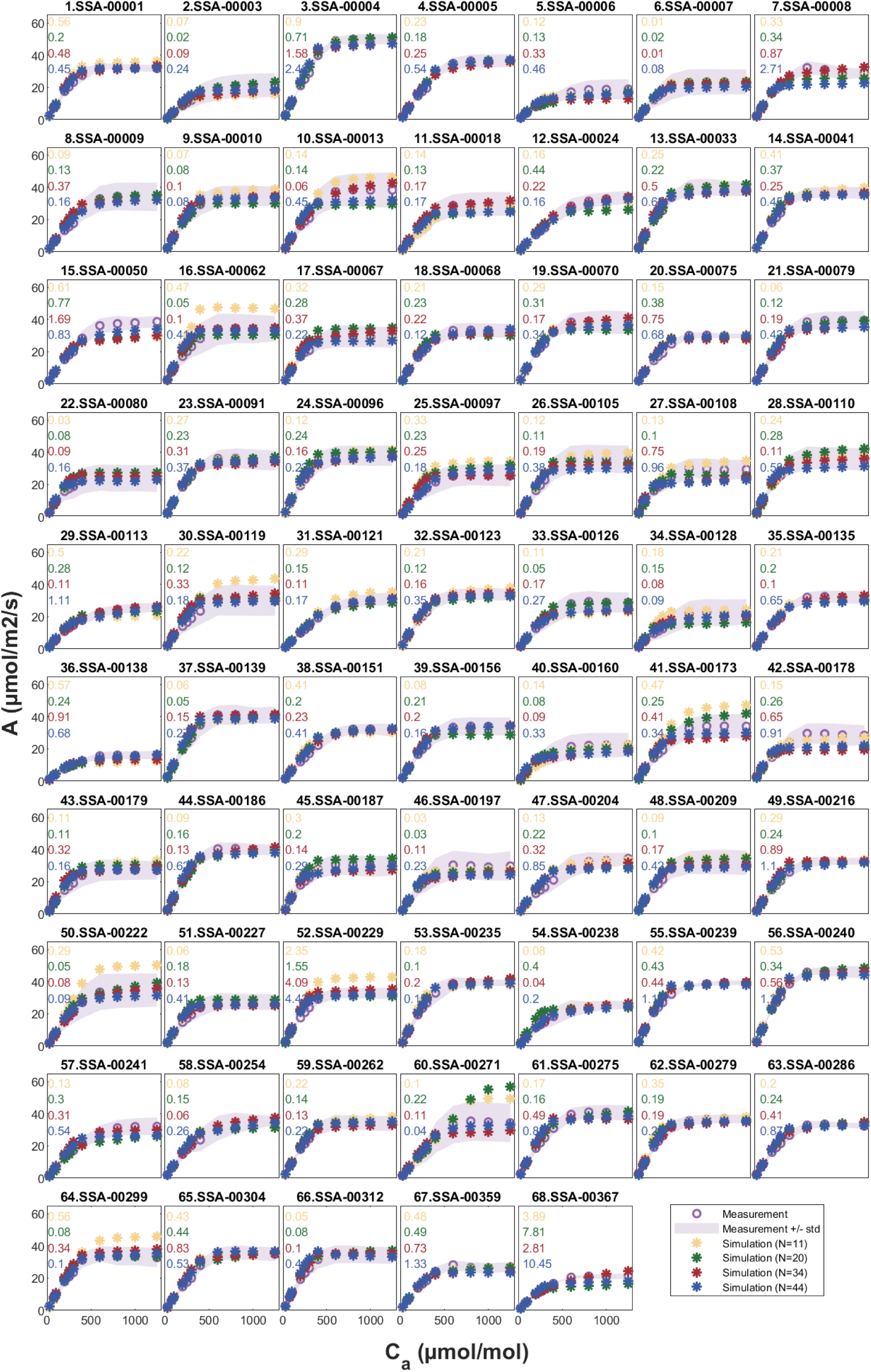
Fitting of photosynthetic rate (*A*) under varying ambient CO_2_ levels (*C_a_*) for 68 genotypes grown in 2023.

**Figure S10:**
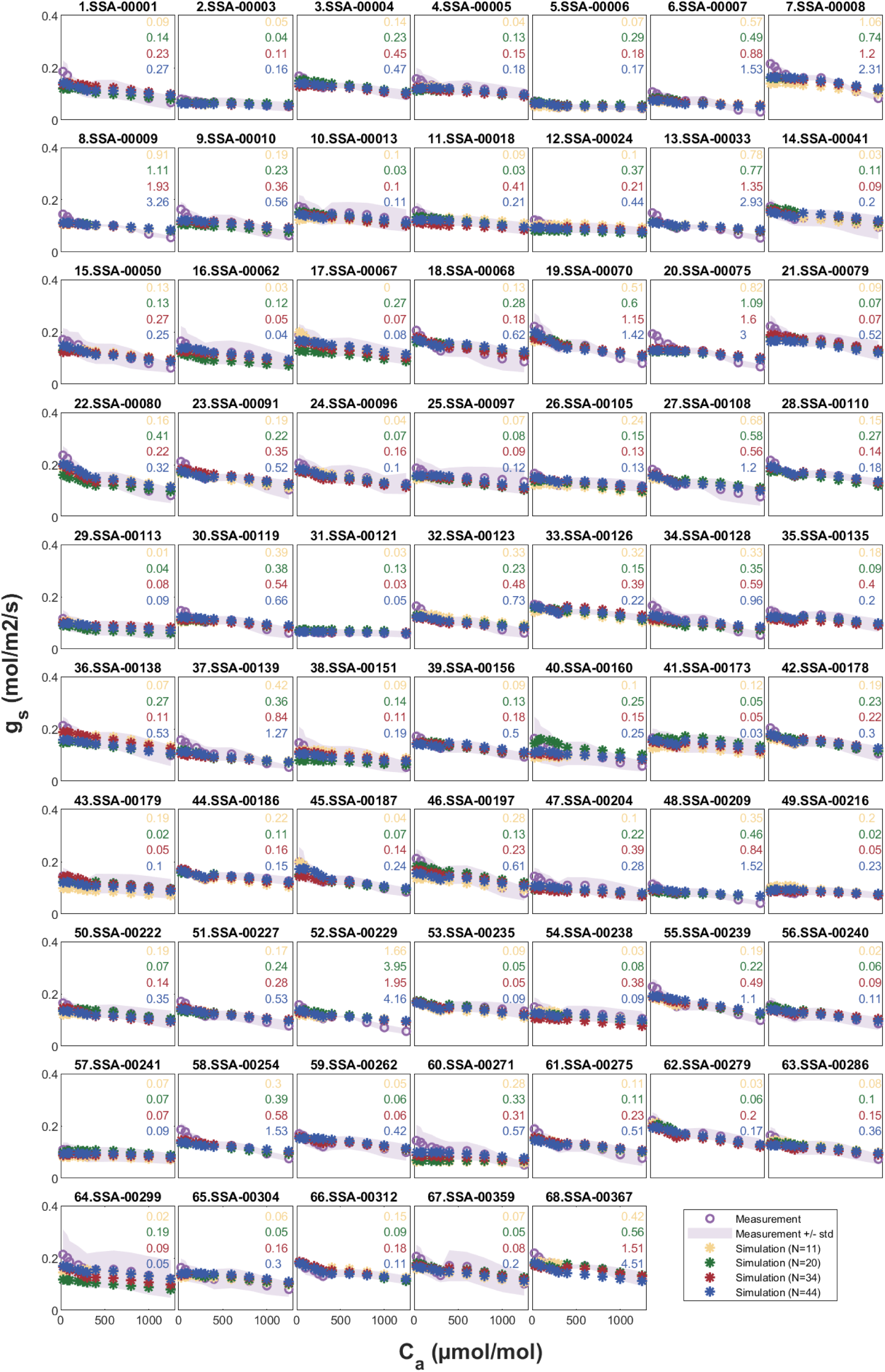
Fitting of stomatal conductance (*g_s_*) under varying ambient CO_2_ levels (*C_a_*) for 68 genotypes grown in 2022.

**Figure S11:**
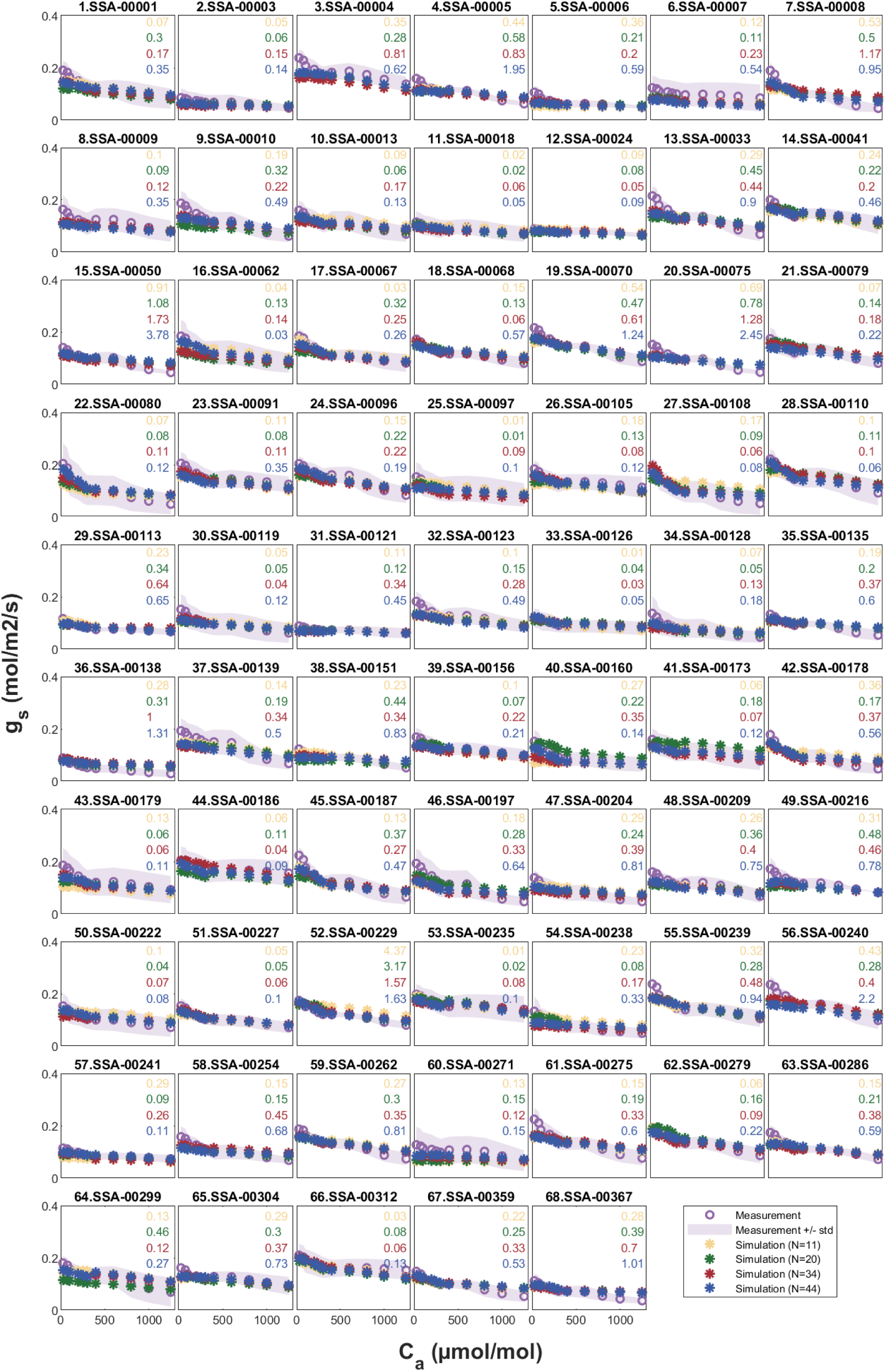
Fitting of stomatal conductance (*g_s_*) under varying ambient CO_2_ levels (*C_a_*) for 68 genotypes grown in 2023.

**Figure S12:**
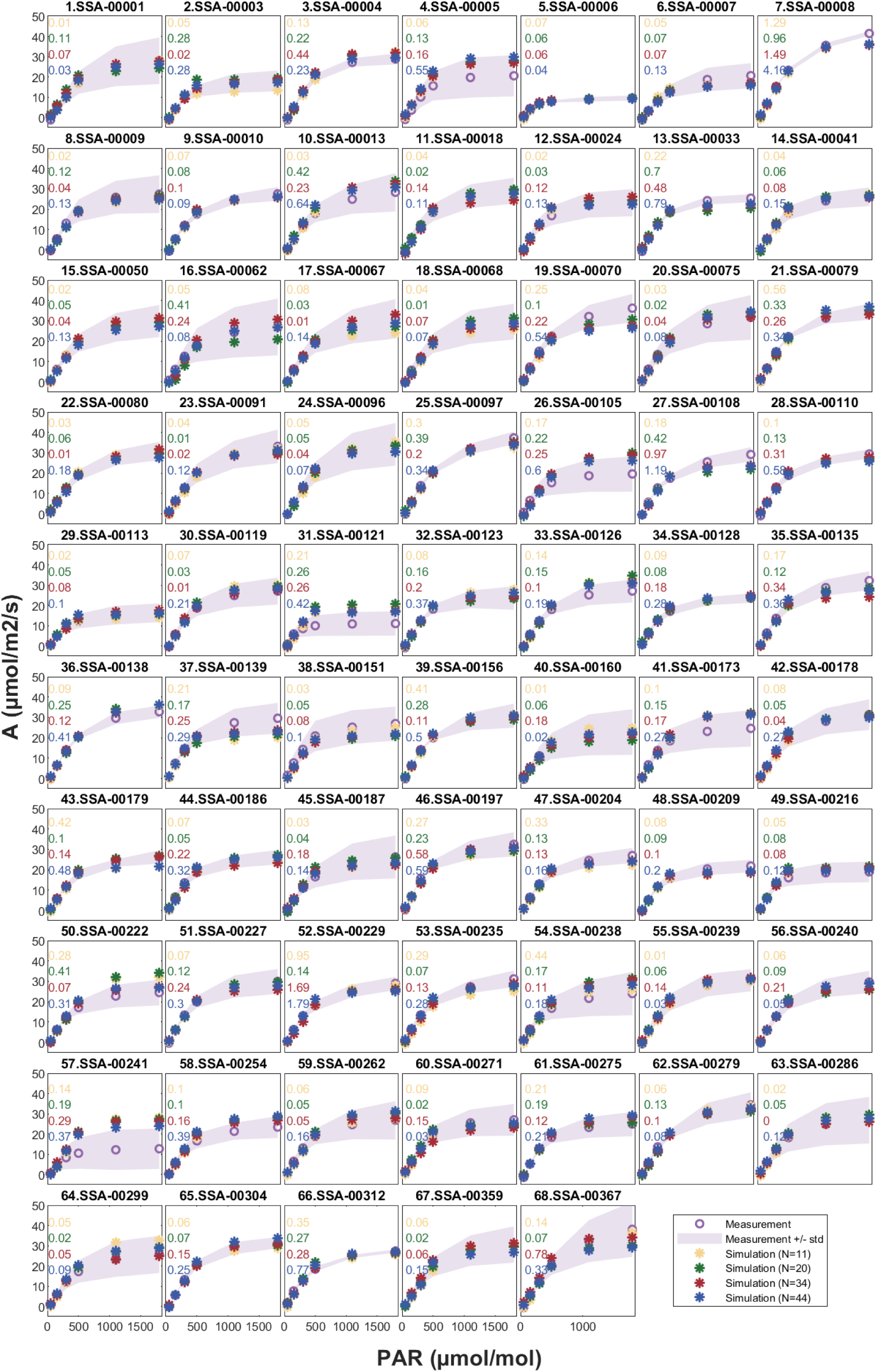
Fitting of photosynthetic rate (*A*) under varying photosynthetically active radiation levels (*PAR*) for 68 genotypes grown in 2022.

**Figure S13:**
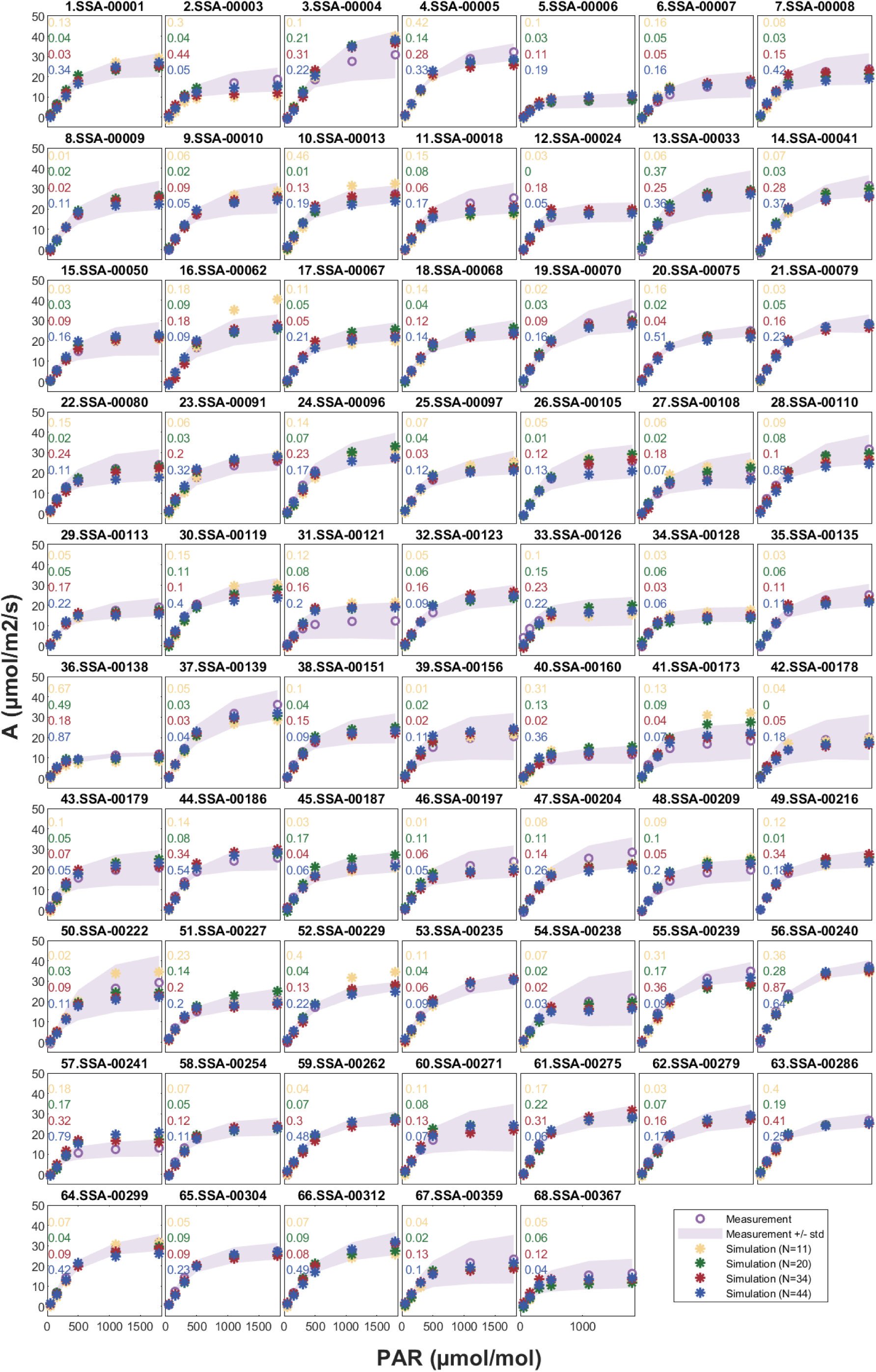
Fitting of photosynthetic rate (*A*) under varying photosynthetically active radiation levels (*PAR*) for 68 genotypes grown in 2023.

**Figure S14:**
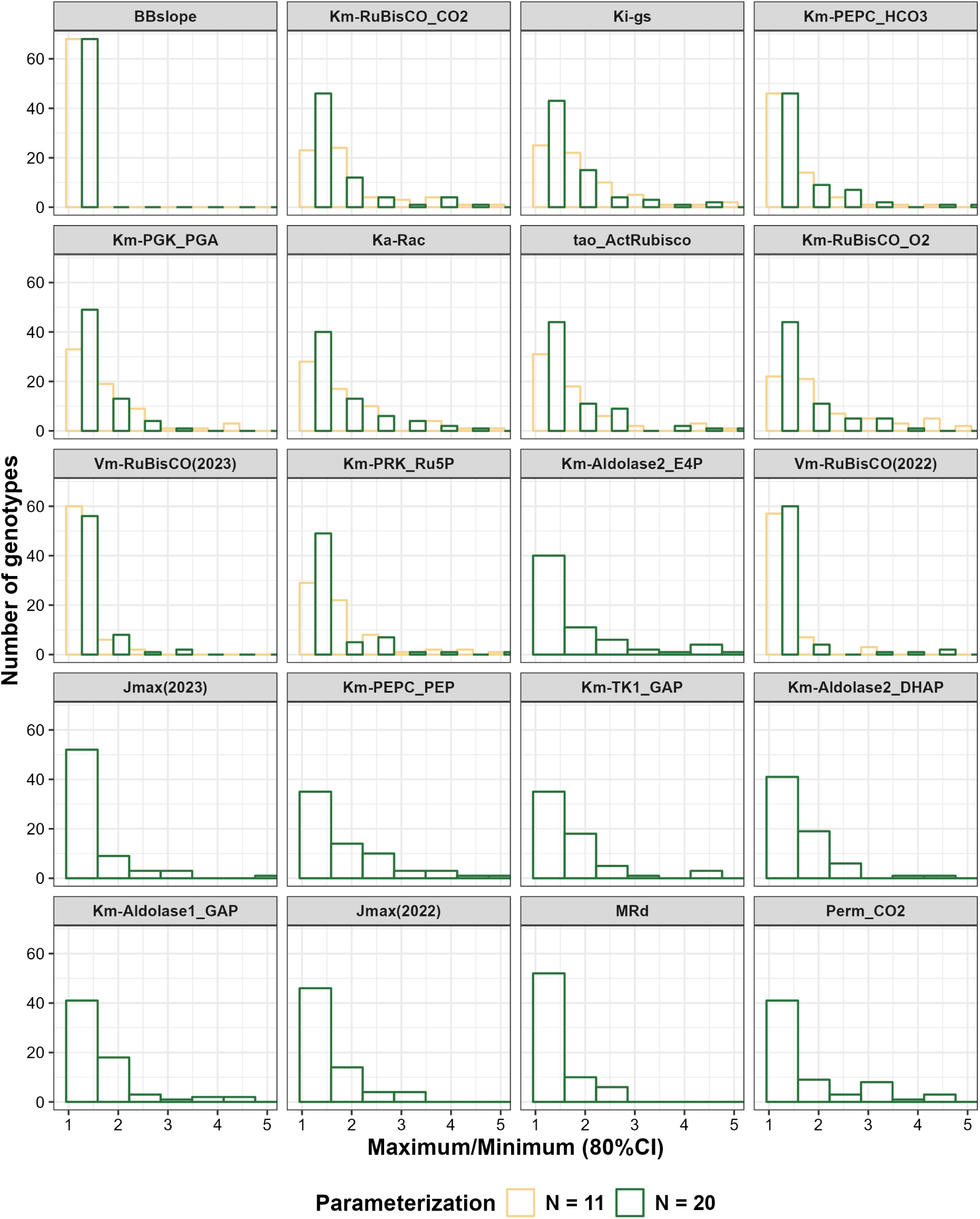
Distribution of fold changes between the upper- and lower-bound for the 80% confidence interval of estimated kinetic parameters across genotypes.

**Figure S15:**
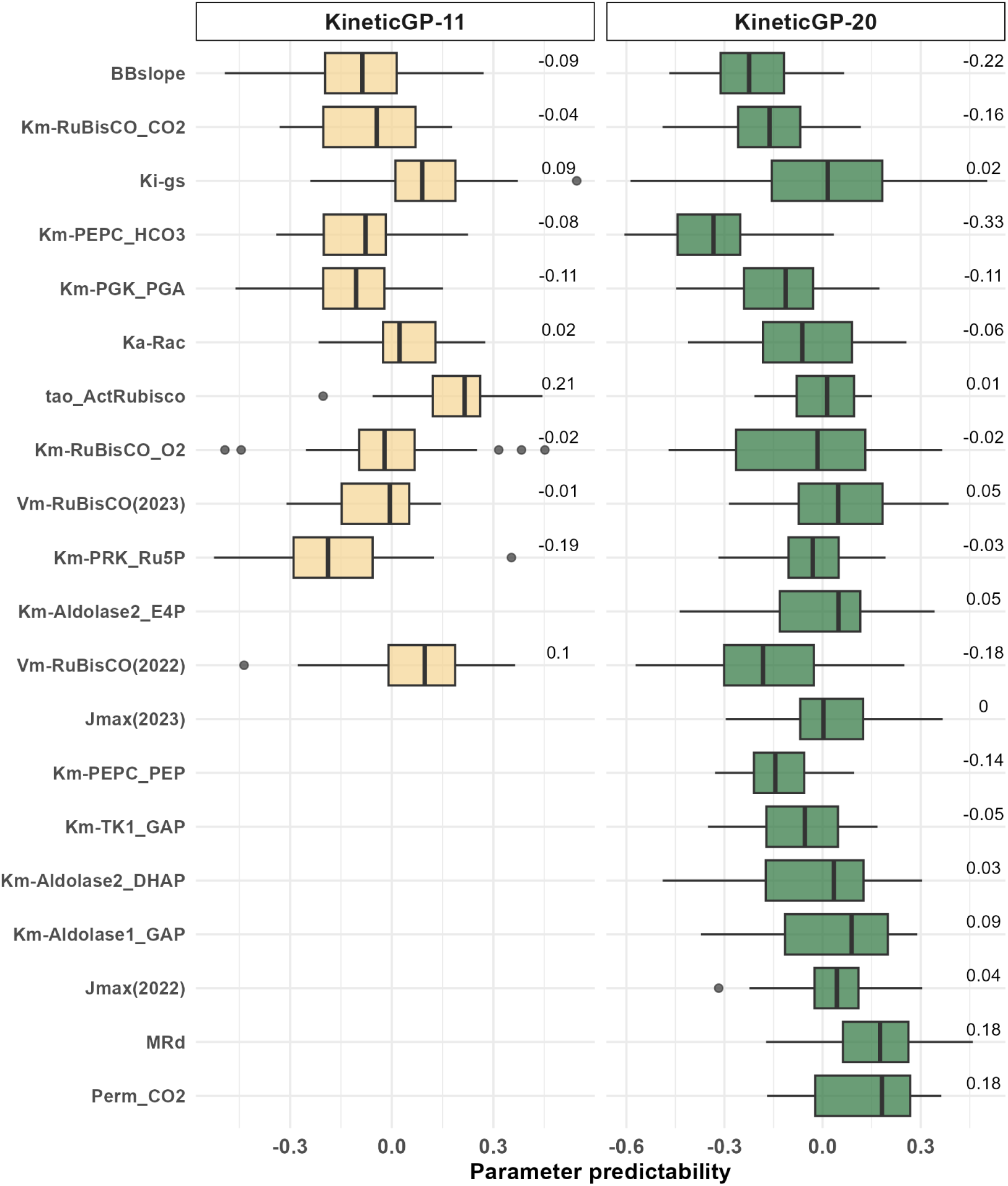
Genomic prediction of kinetic parameters using rrBLUP. Ten repetitions of 3-fold cross-validation were performed to randomly partition the 68 genotypes into training and testing sets. The box-plots show the prediction accuracy across 30 iterations (with median predictability annotated), comparing GP-predicted kinetic parameters from SNP markers with the estimated parameters.

**Figure S16:**
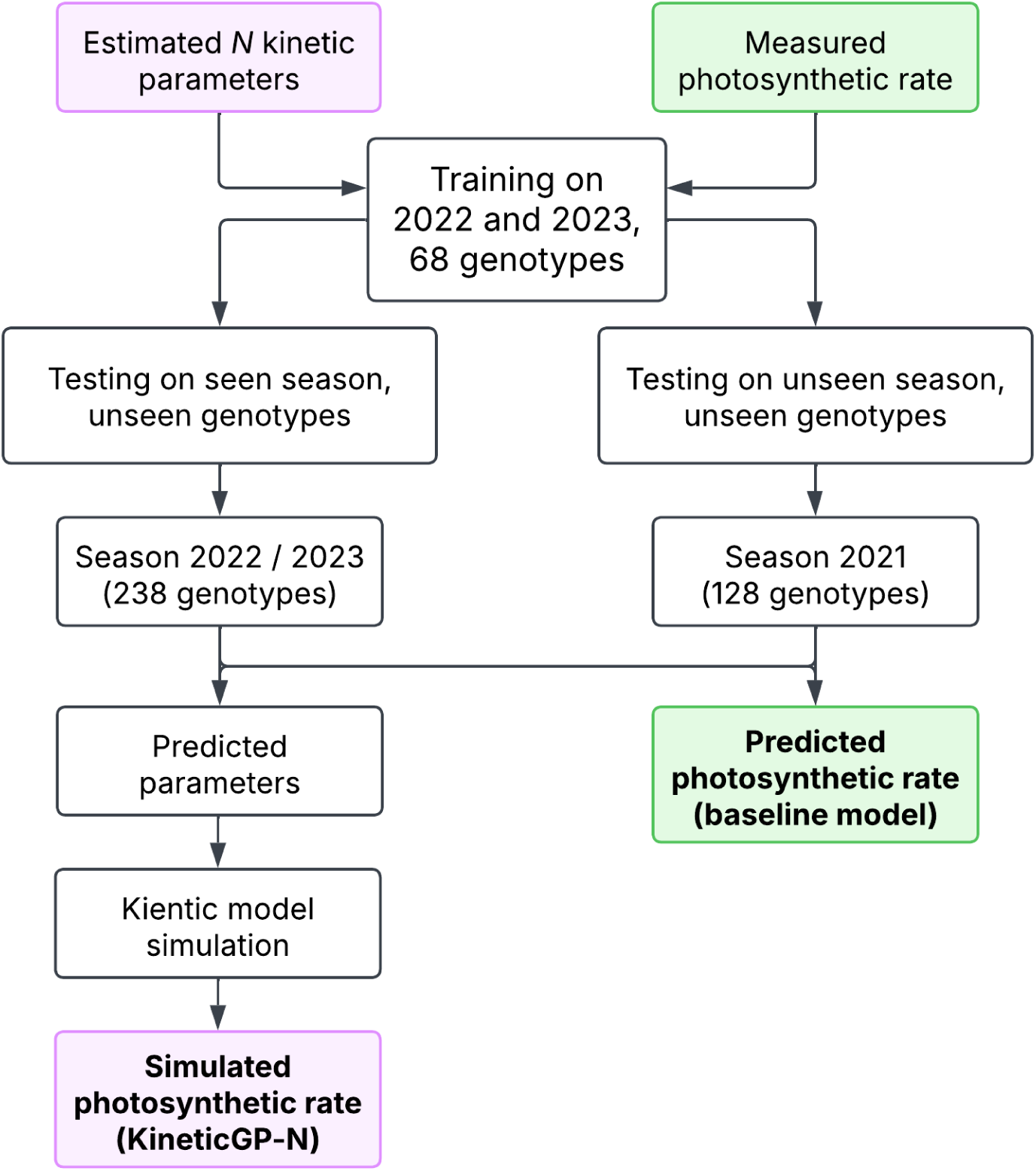
Three different scenarios for testing the predictability of KineticGP in comparison to the baseline model. KineticGP relies on predicting kinetic parameters by training model that use genetic markers as feature and the *N* estimated parameters across genotypes as responses. The predicted parameters for unseen genotypes were used to simulate photosynthesis rate under controlled gas exchange measurement setting (with the same temperature for all measured genotypes). The baseline model uses genomic prediction methods to predict photosynthesis rate directly from measured values.

**Figure S17:**
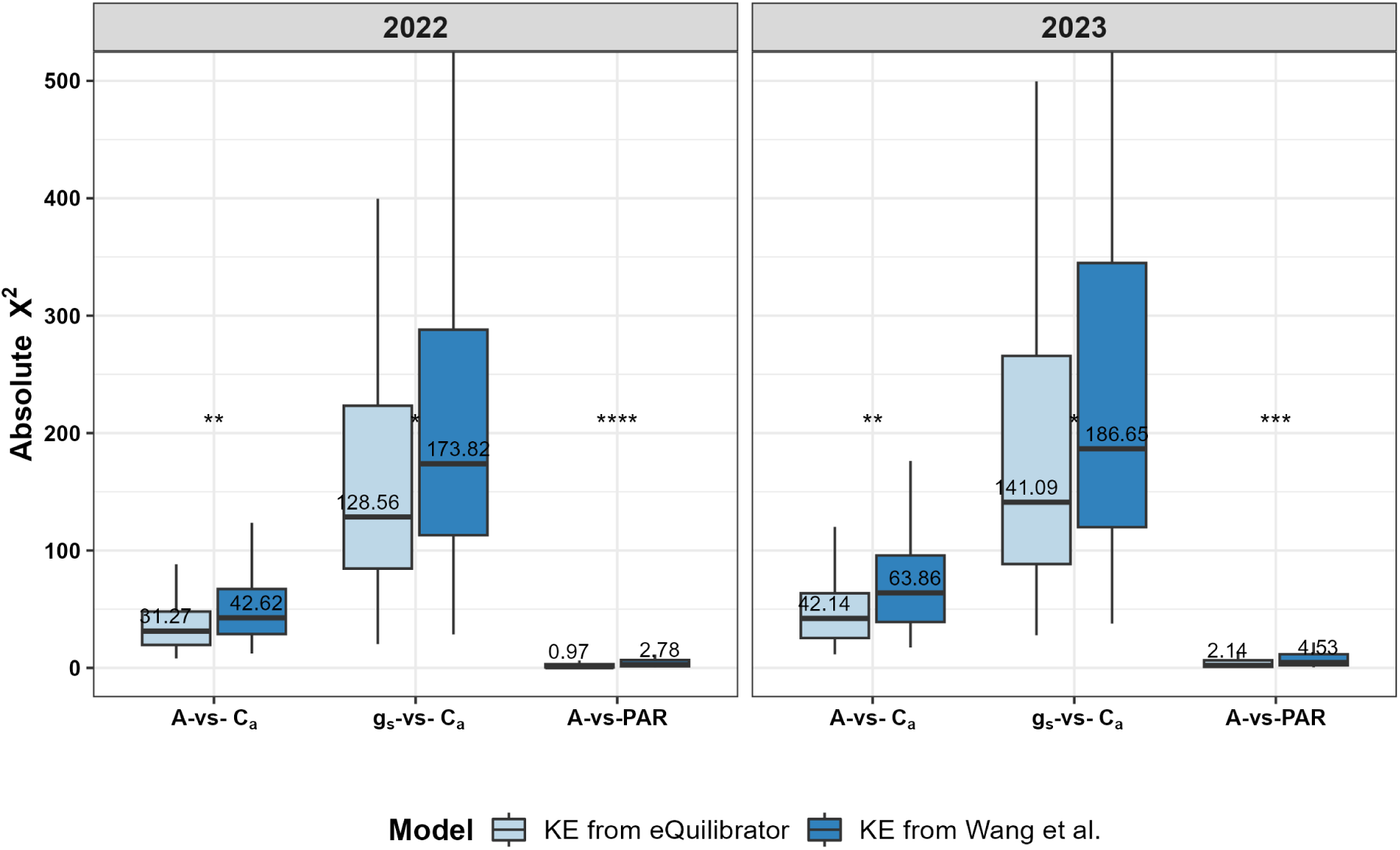
Comparison of initial solution fit between kinetic model using equilibrium constants (K_eq_) from Wang et al. (2021) and from eQuilibrator database. Boxplot of absolute *χ*^2^ statistics of different curves across all 68 genotypes, with different colors representing used models.

**Figure S18:**
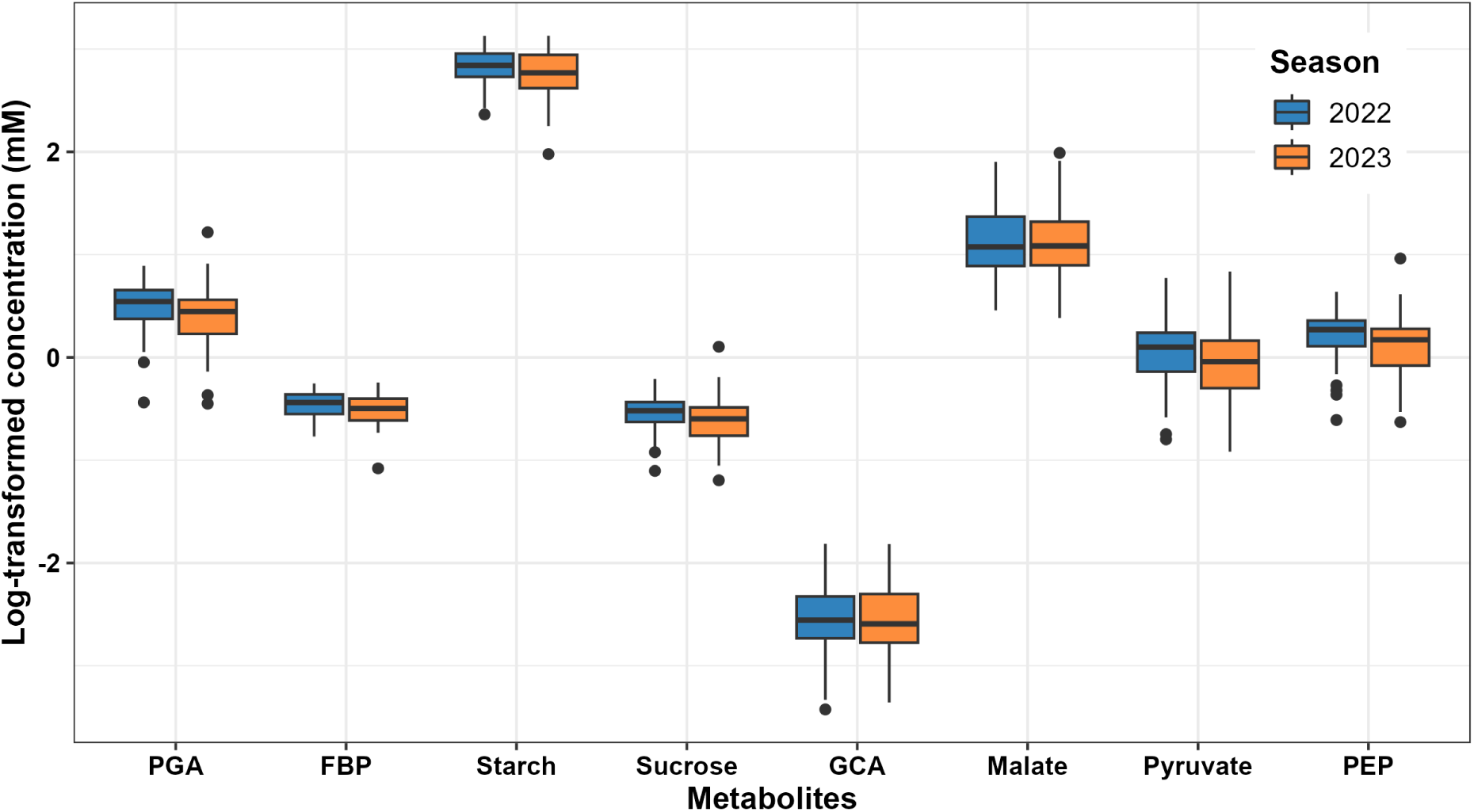
Log-transformed concentration of eight key metabolites in C_4_ photosynthesis after 120 seconds of simulation at 400 µmol CO_2_ mol^-1^ to reach steady state for 68 genotypes using KineticGP-11. The concentration of each metabolite was calculated by summing up the concentration from different compartments.

